# Multi-Modal Kinome Profiling Discovers Mesenchymal-Like Polarity Networks that Underly Directed Hepatocellular Carcinoma Cell Migration

**DOI:** 10.64898/2026.09.11.751090

**Authors:** Thankhoe A. Rants’o, Kathryn Woods, Paige Jensen, Katie A. Walker, Alexandria M. Chan, Kathleen M. Maguire, Rebecca G. Zitnay, Lotfa H. Lovely, Jingshu Yang, Augustine Takyi, Paul Stewart, Kimberley Evason, Robert L. Judson-Torres, Martin Golkowski

**Affiliations:** Department of Pharmacology and Toxicology, University of Utah, Salt Lake City, UT 84112, USA; Huntsman Cancer Institute, University of Utah, Salt Lake City, UT 84112, USA; Department of Oncological Sciences, University of Utah, Salt Lake City, UT 84112, USA; Department of Nutrition and Integrative Physiology, University of Utah, Salt Lake City, UT 84112, USA; Department of Pathology, University of Utah, Salt Lake City, UT 84112, USA; Department of Dermatology, University of Utah, Salt Lake City, UT 84112, USA

**Keywords:** Mass spectrometry, proteomics, protein kinase, hepatocellular carcinoma, epithelial-mesenchymal transition, cell polarity, organelle positioning, DAPK3, FILIP1L

## Abstract

Metastasis and associated therapy resistance remain the principal drivers of cancer related death, and there is a pressing need for a deeper mechanistic understanding and anti-metastatic therapies. For patients that suffer from hepatocellular carcinomas (HCCs), which are the most common primary liver cancers, frequent systemic metastasis results in bleak 5-year survival prognoses of only 4%. To metastasize, carcinoma cells must acquire an invasive phenotype, which typically requires switching from an epithelial-like apical-basal polarity to the front-rear polarity of mesenchymal-like cells. Signaling cues that originate in the tumor microenvironment can activate cellular morphogenic programs that drive polarity switching, like the epithelial-mesenchymal transition (EMT). Protein kinases control most cell signaling pathways and are highly actionable drug targets; however, systematic studies determining the kinases that underly the epithelial-mesenchymal polarity switch (EMPS) are lacking. We developed an assay platform that integrates mass spectrometry (MS)-based kinome profiling, broadly capturing kinase network activity, with chemical genetic screening using selective kinase inhibitors and quantitative phase imaging (QPI), serving as the phenotypic readout. Applying this approach that we dubbed ‘morphokin-MS’, to epithelial-like HCC cell lines that we induced to undergo EMPS identified a conserved network of 12 kinases that contributed to HCC cell polarity switching and directed cell migration; MS-based kinome profiling of 17 HCC patient tumors showed that these kinase are frequently upregulated in human tumors. morphokin-MS also revealed that death associated protein kinase 3 (DAPK3) is one of the principal drivers of the EMPS and directed HCC cell migration. Thus, our mechanistic studies revealed that DAPK3 forms a complex with DAPK1 and filamin-A interacting protein 1-like (FILIP1L), which act as scaffold proteins that recruit DAPK3 to the centrosome. Pharmacological and genetic inhibition of the DAPK1-DAPK3-FILIP1L complex blocked centrosome repositioning and microtubule polarization toward the leading edge of mesenchymal-like HCC cells, directed cell migration, and invasion. Our morphokin-MS method and comprehensive kinome profiling data will serve as a valuable resource for the cancer research community; our discovery of an inducible mesenchymal-like DAPK1-DAPK3-FILIP1L polarity complex that controls centrosome positioning in motile HCC cells may lead to the development of novel therapeutics for combatting cancer metastasis.

## Introduction

Metastasis and associated therapy resistance remain as the principal drivers of cancer related death, and there is a pressing need for a deeper mechanistic understanding and novel anti-metastatic therapies.^1, 2^ To metastasize, carcinoma cells must acquire a migratory and invasive phenotype, which typically requires switching from an epithelial-like apical-basal polarity to the front-rear polarity of mesenchymal-like cells.^3, 4^ Signaling cues that originate in the tumor microenvironment (TME) activate cellular morphogenic programs that drive polarity switching, like the epithelial-mesenchymal transition (EMT).^3, 5–7^ Polarity programs are broadly altered in cancer, and inhibiting their oncogenic functions can block directed cell migration and reduce therapy resistance, and may thus serve as a rational therapeutic target for preventing metastasis and overcome resistance.^8–11^

The 518 human protein kinases, i.e., the kinome, control most cellular signaling networks through reversible protein phosphorylation.^12–15^ Kinases phosphorylate ≥75% of all proteins to alter their activity, protein-protein interactions (PPIs), stability, and localization.^16, 17^ Kinases are frequently dysregulated in cancer, and they are highly actionable with mostly synthetic, ATP-competitive, small molecule kinase inhibitors (KIs).^14, 18–20^ Thus, kinases have emerged as critical cancer drug targets, and 94 KIs have been approved by the US-FDA for the treatment of various cancers and other conditions. Kinases are critical regulators of cell polarity; thus, atypical protein kinase C isoforms (aPKCs) and PAR-1/MARK kinases are well known to control polarity complexes, including Par (Par3/Par6/aPKC), Crumbs (Crumbs/Pals1/PATJ), and Scribble (Scrib/Lgl/Dlg).^3, 21, 22^ The roles of Par, Crumbs, and Scribble are relatively well understood, and they typically promote the apical-basal polarity of epithelial cells, often acting as tumor suppressors.^4, 23–26^ Mesenchymal-specific polarity complexes that may be inducible by TME-derived signals, in contrast, are far less well defined, and systematic studies profiling kinome alterations during epithelial-mesenchymal polarity switching (EMPS) are lacking; this likely left numerous kinases that control oncogenic polarity switching undiscovered.

Hepatocellular carcinoma (HCC) is the most common form of primary liver cancer, and affected ∼1 million new patients in 2025, worldwide.^27–29^ HCC remains a stubborn problem in the clinic due to frequent intrahepatic end extrahepatic metastasis with portal vein thrombosis, rendering surgery and systemic therapies ineffective, and reducing HCC patient’s 5-year survival prospects to a dismal 4%.^30, 31^ In the US and other developed countries, HCC incidence is rapidly increasing due to the obesity epidemic and prevalent metabolic disorders, like type II diabetes, that drive the metabolic dysfunction-associated steatohepatitis (MASH)-HCC disease etiology.^32^ EMT-like phenotypic transitions have been shown to be critical for HCC metastasis and therapy resistance, and recent single cell RNA sequencing (scRNA-seq) studies have confirmed that most HCCs contain cancer cell populations that have undergone such transitions.^30, 33, 34^ Concordantly, loss of cell polarity has been shown to promote HCC initiation and progression, and inhibiting polarity loss can reduce metastasis.^10, 35, 36^ Thus, the EMPS likely promotes HCC metastasis and therapy resistance. Actionable, kinase-mediated signaling pathways that may underly the HCC cell EMPS remain largely obscure, hindering mechanistic insight and developing polarity-targeting therapeutics.

To overcome this challenge, we developed a multimodal assay platform dubbed ‘morphokin-MS’ that integrates our mass spectrometry (MS)-based kinome profiling platform with chemical genetic screening utilizing selective KIs and high-content quantitative phase contrast imaging (QPI).^37–45^ Using this approach, we discovered a kinase network that was frequently induced during HCC cell EMT-like transitions, and that controlled different aspects of the EMPS, including directed cell migration, morphology changes, and organelle positioning. Specifically, we induced four distinct HCC cell lines to undergo the EMPS *in vitro* using a combination of cytokines (TGFβ and WNT5α) and inhibitory antibodies (anti E-cadherin, Dkk-1, and Fsrp-1), and quantified kinome reprogramming using our kinobead AP-MS method.^37^ Kinome reprogramming was highly heterogeneous among cell lines, however, a conserved network of 12 kinases emerged that was consistently activated in 3/4 HCC lines; chemical genetic interrogation of these kinases using QPI and wound healing assays identified death-associated protein kinase 3 (DAPK3) as one of the principal kinases controlling the HCC cell EMPS and directed cell migration. Kinobead AP-MS profiling of 17 HCC patient tumors showed that these kinases are frequently overexpressed in tumors, including DAPK3. Our subsequent mechanistic studies revealed that DAPK3 controls centrosome positioning and microtubule network polarization during directed mesenchymal-like cell migration in a protein phosphorylation-dependent manner. Furthermore, we found that filamin A-interacting protein 1-like (FILIP1L) and DAPK1 are critical scaffold proteins that recruited DAPK3 to the centrosome. Disrupting DAPK1-DAPK3-FILIP1L complex function using siRNAi or dual-specificity pharmacological inhibitors of DAPK1 and DAPK3 blocked HCC cell migration and invasion *in vitro*.

Collectively, we developed the morphokin-MS assay platform that broadly links kinome reprogramming to changes in cell morphology, polarity, and migration. Applying morphokin-MS to HCC cells uncovered a network of inducible mesenchymal-like polarity kinases that may be targeted to prevent EMPS and metastasis. Because cellular morphogenic programs like the EMT are highly conserved across different types of solid cancers and are essential for physiological processes like embryonic development and wound healing, our comprehensive kinome data and approach will broadly serve as a resource for the cell biology and cancer research communities.

## RESULTS

### 1. Functionally interrogating EMT-like transitions and polarity switching *in vitro* with morphokin-MS

To systematically identify kinases that control the EMPS, we designed a multimodal, integrated approach (morphokin-MS) that combines ***1)*** unbiased and quantitative MS-based kinome profiling to identify and prioritize kinases that become induced during EMT-like transitions, and ***2)*** chemical genetic screening with selective KIs and QPI to distinguish kinases that promote the mesenchymal-like polarity from bystanders (**Fig. 1A**). To account for the heterogeneity of EMT signaling, we used four different epithelial-like HCC cell lines, i.e., HuH-7, HepG2, PLC/PRF/5, and Hep3B2.1-7.^39^ First, we validated that we could induce epithelial-like HCC lines to undergo EMT-like transitions and polarity switching, and that we could quantify mesenchymal-like features using our assay platform. We treated HuH-7 cells with an EMT inducing media supplement (hereafter referred to as ‘EMT mix’) composed of the cytokines TGF-β and WNT5α, and antibodies (Abs) that inhibit E-cadherin (CDH1) and the canonical WNT inhibitors secreted frizzled-related protein 1 (sFRP-1) and dickkopf-related protein 1 (Dkk-1) for 4 days, followed by confocal immunofluorescence (IF) imaging of fibrous (F)-actin and N-Cadherin (CDH2; **Fig. 1B**).^46^ IF imaging showed F-actin stress fiber formation and N-cadherin expression at the peripheral membrane, which are markers of a motile, mesenchymal-like phenotype.^47^ Next, we performed QPI using the Phasefocus Livecyte imaging system to determine morphology changes in HuH-7 cells that have been treated with EMT mix for 4 days. Brightfield images showed the switching from the typical cobblestone pattern of epithelial-like HCC cells to an elongated spindle-shaped morphology (**Fig. 1C**). Quantifying HuH-7 cell morphology showed significantly reduced sphericity (roundness) and increased cell area, which was concordant with a more mesenchymal-like morphology (**Fig. 1D**). Random cell migration measured by Livecyte imaging and directed cell migration determined by trans-well migration assay were also significantly increased, validating the transition to a more motile phenotype, and that QPI can capture mesenchymal-like features (**Fig. 1E** and **1F**). Next, to more systematically validate that we can induce EMT-like transitions, we performed quantitative MS-based global proteome profiling of HuH-7, HepG2, PLC/PRF/5, and Hep3B2.1-7 cells that we treated with EMT-mix for 4 days; this quantified a total of 10,961 proteins in the four cell lines (**Table S1**). Differential expression analysis (DEA) comparing vehicle controls with EMT mix treatment showed consistently increased expression of the EMT-inducing transcription factors SMAD3, JUNB,^48^ STAT3,^49^ and SOX4,^50^ and the SMAD3 transcriptional targets PDGFA and TGFBI, as well as the stress fiber-forming vimentin (VIM), and different collagens (**Fig. 1G** and **Table S1**). Transferrin (TF) and albumin (ALB) decreased in expression, as did other markers of hepatocyte differentiation like the transcription factor HNF1A (**Fig. 1G**). Concordant with EMT-induced cell cycle arrest, the cell cycle inhibitor p14-INK4b (CDNK2B) was increased, while the epithelial stem cell marker GPC3 and the proliferation marker MYC were decreased (**Fig. 1G**). Reduced proliferation was also evident from Livecyte imaging, which showed two-fold prolonged doubling times in EMT mix treated HuH-7 cells (**Fig. S1A**). Immunoblotting validated the trend for increased expression of the mesenchymal markers VIM, FN1, SMAD3 and JUNB, and the decreased expression of hepatocyte markers TF and ALB in response to the EMT mix (**Fig. 1H** and **S1B**). Applying gene set enrichment analysis (GSEA) with gene ontology – biological process (GOBP) terms to our global proteomics data also validated the systematic activation of pathways that control cell motility, morphology, polarity and survival (RTK, integrin, and PI3K/AKT signaling), as well as stress responses (Nf-κB and JNK signaling) and development (EMT, TGFβ, WNT, and NOTCH signaling), and the decreased activity of hepatic metabolic pathways (**Fig. 1I** and **Table S1**). Collectively, our imaging and proteomics data demonstrated that we can reliably induce EMT-like transitions and the EMPS in HCC cells *in vitro* and that QPI can quantify feature changes associated with the EMPS.

**Figure 1.**
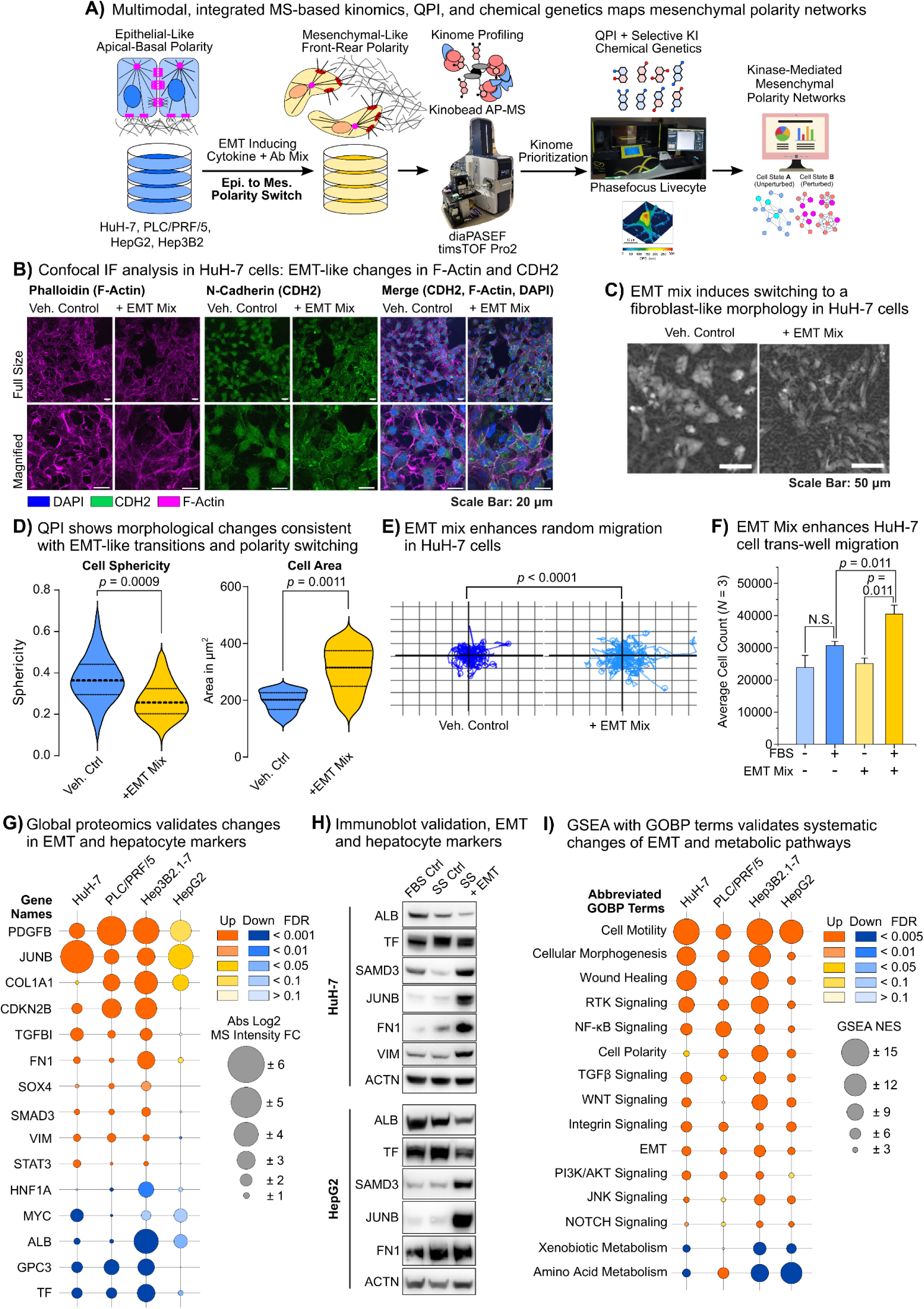
Workflow overview of our morphokin-MS assay platform and method and model system validation. **(A)** Schematic of the morphokin-MS assay platform, including MS-based kinome profiling, chemical genetic screening, and QPI imaging. **(B)** Confocal IF imaging of nucleus (DAPI), F-actin (Phalloidin), and N-cadherin (CDH2, Ab) in HuH-7 cells that were treated with EMT mix in serum-free medium compared to serum-free medium alone for 4 days. **(C)** Bright field images taken on the Phasefocus Livecyte imaging system showing HuH-7 cells subjected to the same treatment as in (B). **(D)** Random cell migration of HuH-7 cells subjected to the same treatment as in (B) captured using the Phasefocus Livecyte imaging system. Experiments were done in technical triplicates and on average 215 cells were imaged in each experiment. *Statistics*: two-sample Student’s T-test, p < 0.05. **(E)** Differences in roundness (sphericity) and cell area of HuH-7 cells subjected to the same treatment as in (B) captured using the Phasefocus Livecyte imaging system. Experiments were done in technical triplicates and on average 215 cells were imaged in each experiment. *Statistics*: two-sample Student’s T-test, p < 0.05. **(F)** Trans-well migration assay of HuH-7 cells treated like in (B) using FBS as the attractant. Error bars are the standard deviation (S.D.). *Statistics*: two-sided two-sample Student’s T-test, p < 0.05. **(G)** MS-based global proteomic analysis of HuH-7 cells treated like in (B), comparing 4-day EMT mix vs. serum-free medium alone. Refers to **Table S1**. *Statistics*: two-sided two-sample Student’s T-test with Benjamini-Hochberg (BH) correction for multiple hypothesis testing, global FDR = 0.05. **(H)** Immunoblot validation of specific EMT markers in HuH-7 and HepG2 cells treated with vehicle in serum-containing medium (FBS ctrl), with vehicle in serum-free medium (serum starvation, SS), and EMT mix in serum-free medium (SS). Relates to **Fig. S1B**. **(I)** GSEA results from analyzing the global proteomics data form (G), showing selected EMT and polarity associated pathway terms. Relates to **Table S1**.

### 2. morphokin-MS identifies a conserved kinase network underlying the HCC cell EMPS

Next, to systematically quantify kinome reprogramming during EMT-like transitions, we applied our kinobead AP-MS protocol to HuH-7, HepG2, PLC/PRF/5, and Hep3B2.1-7 cells that we treated with EMT mix for 4 days;^37^ this quantified 372 protein kinases (**Table S1**). Comparing differentially expressed kinases following EMT mix treatment between the four HCC lines revealed the extensive heterogeneity of kinome reprogramming (**Fig. 2A**). Thus, in each cell line on average 46 (35 – 54; total of 131) and 68 (19 – 167; total of 185) kinases either increased or decreased in abundance; however, compared to the total number of increased or decreased kinases only 9.2% (*N* = 12) and 7.0% (*N* = 13) were significantly increased or decreased in at least 3/4 HCC lines, respectively (**Fig. 2A**). The conserved module of 12 kinases that increased in abundance during the EMT-like transitions in 3/4 HCC lines included the receptor kinases (RTKs) ephrin type-B receptor 2 (EPHB2) and epithelial discoidin domain-containing receptor 1 (DDR1), and the bone morphogenetic protein receptor type-2 (BMPR2), and a diverse set of non-receptor kinases, i.e., NUAK family SNF1-like kinase 1 (NUAK1), death-associated protein kinase 3 (DAPK3), mitogen-activated protein kinase kinase kinase kinase 4 (MAP4K4), protein kinase lysine-deficient 1 (WNK1), protein kinase C delta type (PKCδ or PRKCD), cyclin-dependent kinase 19 (CDK19), the lymphocyte-oriented kinase (LOK or STK10), and AMPK subunit alpha-2 (PRKAA2), as well as the catalytically inactive kinase suppressor of Ras 1 (KSR1, **Fig. 2B**). Rationalizing that kinases that change in abundance across multiple different HCC model cell lines may have conserved functions in the EMPS, we prioritized kinases for our follow-up experiments based on the frequency with which they were significantly increased (≥3/4 cell lines, **Table S1**). Immunoblotting validated the increased abundance of most kinases in EMT mix treated HCC cells, except for WNK1, PRKCD, CDK19, PRKAA2, and KSR1 (**Fig. 2C** and **Fig. S1C**). Additionally, reanalyzing our kinobead AP-MS data from 17 HCC cell lines that stably expressed either an epithelial-like phenotype (*N* = 7) or mesenchymal-like phenotype (*N* = 10) showed that EPHB2, NUAK1, DAPK3, BMPR2, DDR1, and STK10 were also significantly enriched in stably mesenchymal-like lines, suggesting that these kinases are broadly associated with mesenchymal-like states in HCC cells and important for long-term maintenance of mesenchymal-like features (**Fig. S1D**).^39^ Next, to determine which of the EMT mix induced kinases control cell morphology, we performed a chemical genetic screen, treating the stably mesenchymal-like HCC line SNU387 with one or two selective KIs targeting each of the seven validated kinases that increased in response to the EMT mix; previous results from our lab showed that SNU387 cells express these kinases at high levels.^39^ We performed QPI analysis using cell sphericity as the readout for changes in cell morphology (**Fig. 2D**). This showed that DAPK3, DDR1, EPHB2, and dual NUAK1 and NUAK2 inhibition significantly increased sphericity, indicating a shift to a more epithelial-like morphology; the Srcfamiliy kinase (SFK) inhibitor Saracatinib served as the positive control, as SFKs are well known to promote a mesenchymal-like morphology.^51^ Next, we quantified changes in directed cell migration using a wound scratch assay; this showed that DAPK3 inhibition completely blocked wound closure in SNU387 cells, and that MAP4K4 and EPHB2 inhibition partially blocked wound closure, as did the positive control Saracatinib (**Fig. 2E**); the other KIs did not significantly reduce wound closure. We validated that DAPK3 inhibition also blocked wound closure in the mesenchymal-like HCC line SNU449 (**Fig. S1E**). Supporting that directed cell migration was inhibited, and not proliferation, QPI doubling times were unaffected by all KIs except Saracatinib, and cell viability assays using cellular ATP content as the readout confirmed that DAKP3 inhibition does not affect SNU387 or SNU449 cell viability (**Fig. S1F** and **S1G**). DAPK3 inhibition also significantly slowed SNU387 cell invasion in a trans-well assay (**Fig. S1H**). Collectively, our results suggested that DAPK3 and other kinases that partake in an inducible mesenchymal-like polarity network promote HCC cell motility. Next, to determine if EMPS-related kinases are overexpressed in HCC patient tumors, we applied kinobead AP-MS profiling to 17 paired HCC tumor and non-tumor liver (NTL) tissues that we received from the Huntsman Cancer Institute (HCI) Biorepository and Molecular Pathology (BMP) Shared Resources (**Table S1**). Tissue samples had diverse underlying etiologies, including six cases with MASH-HCC, four with HCV, one each with HBV, ethanol use, and alpha-1 antitrypsin (A1AT) deficiency, and three with non-cirrhotic livers and no obvious comorbidities, respectively. Kinome profiling quantified 399 kinases; differential expression analysis between paired tumors and NTL tissues showed frequent upregulation of kinases involved in cell cycle regulation and DNA damage checkpoint signaling (e.g., AURKA, CHEK1, PRKDC), proliferation (e.g., MAPK1 and MAPK3), redox stress signaling (e.g., VRK2 and OXSR1), vesicle trafficking (e.g., SCYL1), and mitochondrial function (ADCK1; **Table S1** and **Fig. S2A**). Inducible kinases that we found to be associated with mesenchymal-like HCC states *in vitro* were also frequently upregulated in tumors, particularly EPHB2 (*N* = 14), BMPR2 (*N* = 10) and DAPK3 (*N* = 10, **Fig. 2F** and **Fig. S2B**). Mining The Protein Atlas database revealed that that DAPK3 and BMPR2, among others, also significantly correlated with shorter HCC patient survival (**Fig. 2G** and **Fig. S2C**).^52^ Collectively, kinome profiling of human HCC tissues revealed the frequent upregulation of kinases associated with the EMPS; particularly, DAPK3 emerged as one of the principal kinases promoting a mesenchymal-like morphology and HCC cell directed migration and invasion *in vitro*; DAPK3 was also frequently increased in expression and correlated with shorter survival in HCC patients.

**Figure 2.**
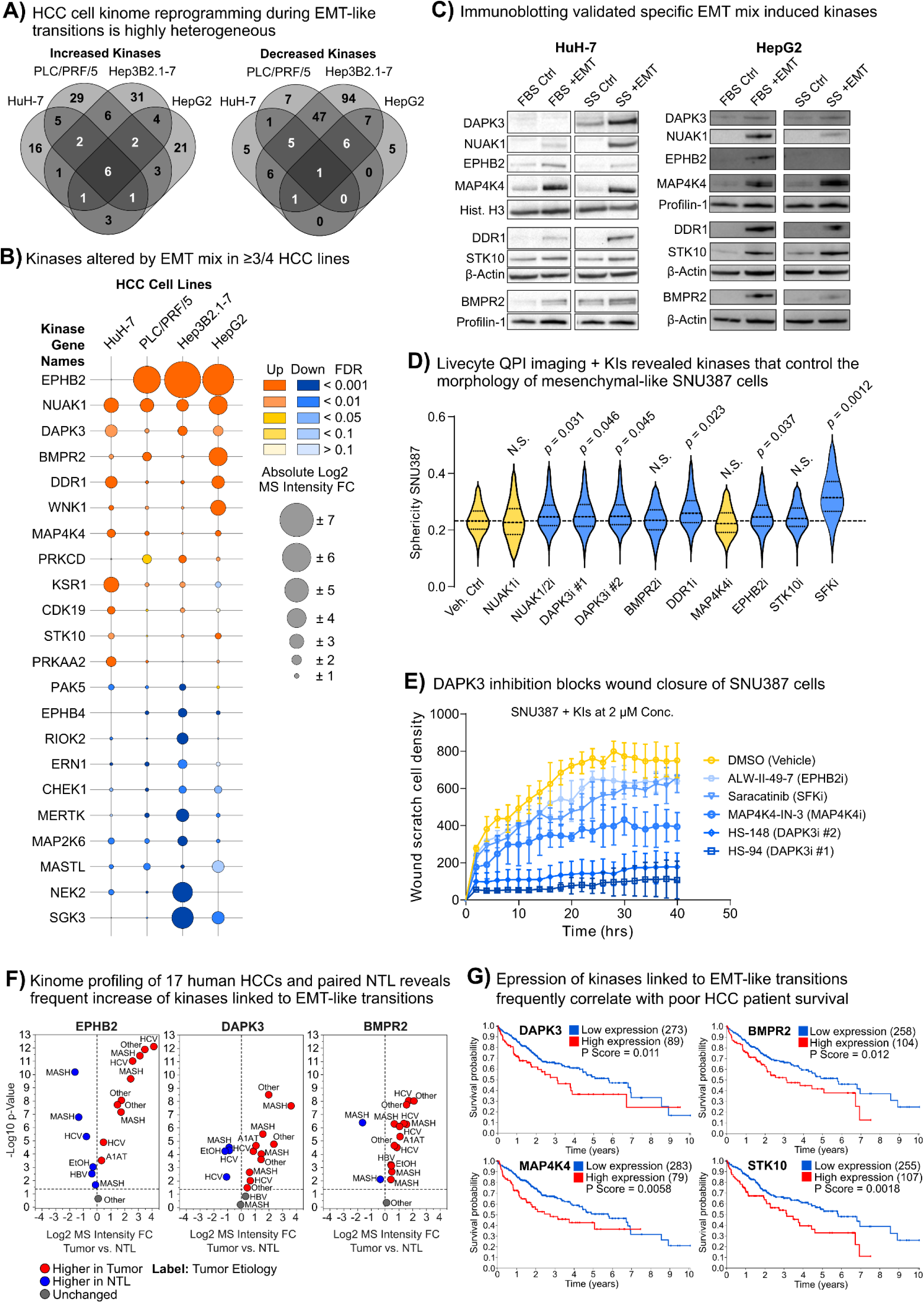
morphokin-MS identifies a conserved network of kinases that promote the HCC cell EMPS and directed cell migration; kinobead AP-MS analysis of human HCC tumors reveals frequent upregulation of several of these kinases *in vivo*. **(A)** Comparing kinases that significantly increased or decreased in abundance in the 4 EMT mix treated HCC cell lines showed extensive heterogeneity in kinome reprogramming. Refers to **Table S1**. *Statistics*: two-sided two-sample T-test with BH correction, global FDR = 0.05. **(B)** Kinobead AP-MS results showing kinases that significantly increased or decreased in abundance in ≥3/4 EMT mix treated HCC cell lines. Refers to **Table S1**. *Statistics*: see (A). **(C)** Immunoblot validation of 7 of the 12 kinases that increased in abundance in response to the EMT mix. Refers to **Fig. S1C**. **(D)** QPI using the Phasefocus Livecyte imaging system of SNU387 cells that were treated with selective KIs of the validated EMPS-associated kinases for 4 days. Increased cell roundness (sphericity) was used as a measure for Kis causing the switching from a mesenchymal-like morphology to an epithelial-like morphology. Saracatinib (SFK inhibitor) was positive control. Experiments were done in technical triplicates and on average 215 cells were imaged in each experiment. *Statistics*: two-sample T-test, p < 0.05. **(E)** 96-well plate wound scratch assay of SNU387 cells pre-treated with KIs targeting the validated EMPS-associated kinases for 72 h; all KI concentrations were 2 µM. Saracatinib (SFK inhibitor) was positive control. Refers to **Fig. S1E**. Error bars are the S.D. **(F)** Kinobead AP-MS profiling of 17 HCC patient tumors and paired non-tumor liver (NTL) tissues shows frequently increased abundance of EMP-associated kinases in tumors compared to NTL tissues. Refers to **Fig. S2A** and **S2B**, and **Table S1**. Statistics: see (A). **(G)** Expression of EMPS-associated kinases broadly correlated with shorter patient survival; source was The Protein Atlas. Refers to **Fig. S2C**.

### 3. FILIP1L and DAPK1 act as scaffold proteins that co-recruit DAPK3 to the centrosome in mesenchymal-like HCC cells

Next, to validate the mechanism of action (MOA) of the DAPK3 inhibitors HS-94 and HS-148, we performed a kinome-wide competitive binding assay using kinobead AP-MS in mesenchymal-like SNU761 cell lysates, an HCC cell line that we found to express very high levels of DAPK3.^37, 39^ This showed that both inhibitors bound several additional kinases, i.e., DAPK1, casein kinase 2 (CK2 or CSNK2A1/2), and the CK2-interacting kinases BRD2, CDK11B, and DYRK1A (**Fig. 3A** and **Table S2**).^37^ This suggested that DAPK1 and CK2 may present high affinity off-targets of HS-94 and HS-148, and that the remaining kinases were co-competed with CK2. To confirm binding, we performed an *in vitro* kinase assay with Reaction Biology, showing that both HS-94 and HS-148 are equipotent inhibitors of DAPK1 and DAPK3 with IC_50_ values of 22.5 to 47.9 nM (**Table S2**). In contrast, HS-94 and HS-148 showed IC_50_ values between 2.88 µM and 6.51 µM for CK2. We concluded that HS-94 and HS-148 are potent and selective dual-specificity inhibitors of the closely related DAPK1 and DAPK3.

**Figure 3.**
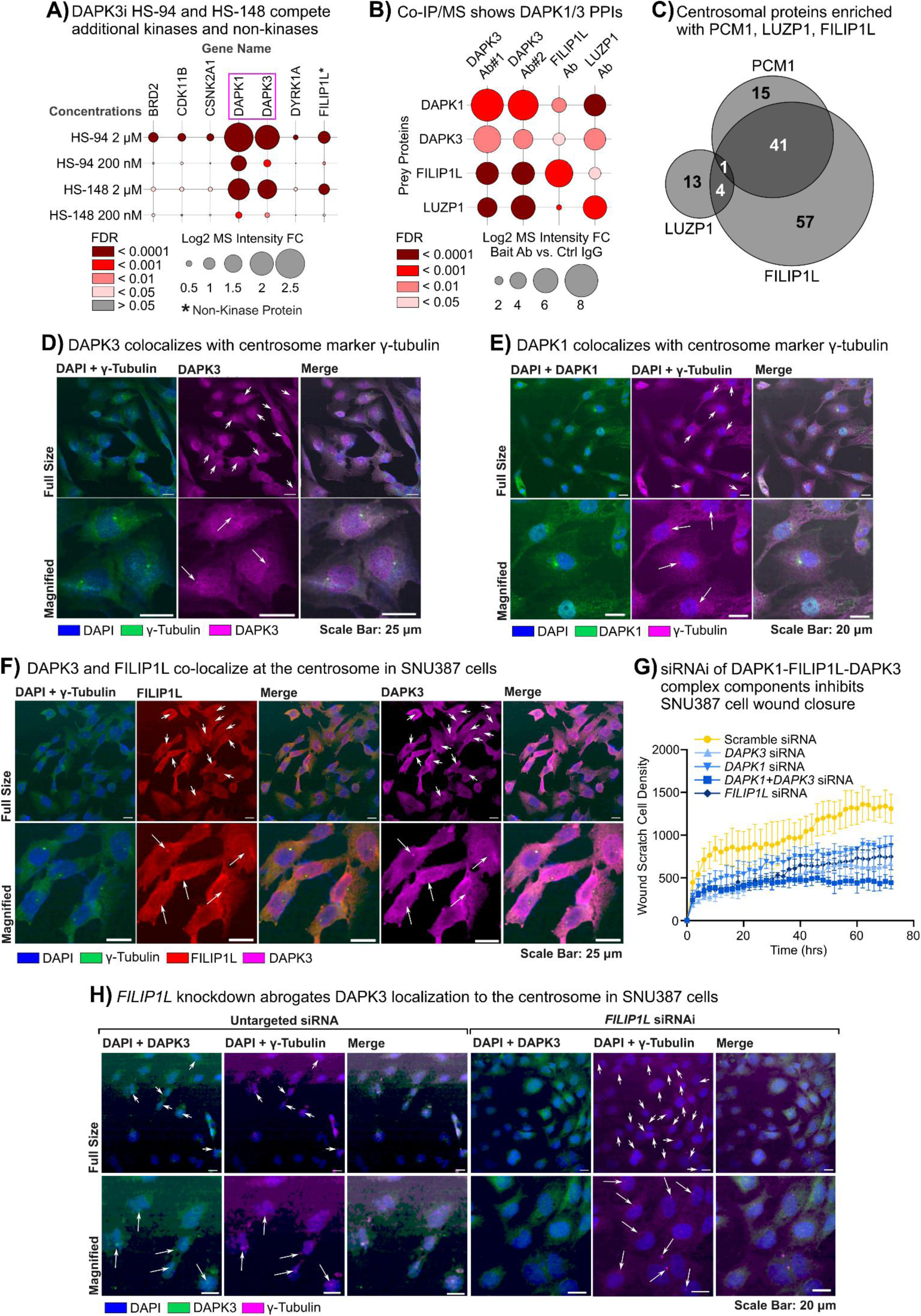
DAPK3 interacts with DAPK1 and FILIP1L at the centrosome. **(A)** Kinome-wide selectivity profiling of the DAPK3 inhibitors HS-94 and HS-148 in SNU761 cell lysates using kinobead AP-MS. Only proteins identified with >2 prototypic peptides are shown. Refers to **Table S2**. *Statistics*: two-sided two-sample T-test with BH correction, global FDR = 0.05. **(B)** Ab-based co-IP/MS of DAPK3, FILIP1L, and LUZP1 in SNU387 and SNU449 cell lysates. Refers to **Table S2**. *Statistics*: two-sided two-sample T-test with BH correction, p < 0.05. **(C)** Comparison of centrosomal proteins (GOCC: ‘Centrosome’) that were co-enriched with Abs targeting FILIP1L, LUZP1, and PCM1 in our co-IP/MS experiments. Refers to **Table S2**. *Statistics*: see (B). **(D)** Co-localization study using confocal IF, staining for the nucleus (DAPI), the centrosome (γ-tubulin), and DAPK3 in SNU387 cells. **(E)** Co-localization study using confocal IF, staining for the nucleus (DAPI), the centrosome (γ-tubulin), and DAPK1 in SNU387 cells. **(F)** Co-localization study using confocal IF, staining for the nucleus (DAPI), the centrosome (γ-tubulin), DAPK3, and FILIP1L in SNU387 cells. **(G)** 96-well wound scratch assay of SNU387 cells pre-treated siRNAs targeting *DAPK1*, *DAPK3*, *FILIP1L*, and *DAP1*-*DAPK3* combined for 3 days. A scrambled, untargeted siRNA sequence was the negative control. Refers to **Fig. S3C** and **S3D**. Error bars are the S.D. **(H)** Co-localization study using confocal IF, staining for the nucleus (DAPI), the centrosome (γ-tubulin), and DAPK3 in SNU387 cells that were pretreated with a FILIP1L siRNA sequence for 3 days. A scrambled, untargeted siRNA sequence was the negative control. Refers to **Fig. S3E**.

Notably, HS-94 and HS-148 also significantly co-competed one non-kinase protein in our kinobead AP-MS assay, the filamin A-interacting protein 1-like (FILIP1L); leucine zipper protein 1 (LUZP1), a known interactor of DAPK3,^53, 54^ was co-competed as well, but did not reach significance (**Fig. 3A** and **Table S2**). Like DAPK3, LUZP1 and FILIP1L contain leucine zipper domains that enable homo- and heterodimerization with other leucine zipper proteins; this suggested that FILIP1L, and possible LUZP1, interacted with DAPK3 in mesenchymal-like HCC cells.^55, 56^ To validate these interactions, we next performed co-IP/MS analyses in mesenchymal-like SNU387 cell lysate using two specific DAPK3 Abs that recognize distinct epitopes, and one Ab each recognizing LUZP1 and FILIP1L (**Fig. 3B, Table S2**). This showed that DAPK3 antibodies co-enriched LUZP1 and FILIP1L, and surprisingly DAPK1. LUZP1 and FILIP1L Abs also co-enriched DAPK1 and DAPK3, as well as each other, albeit to a lesser extent (**Fig. 3B**); this added evidence that DAPK1, DAPK3, LUZP1, and FILIP1L form complexes in mesenchymal-like HCC cells. LUZP1 and FILIP1L were shown to localize to the centrosome previously, which plays a critical role in the EMPS and mesenchymal-like cell migration,^8, 56–58^ and our Co-IP/MS data showed that the FILIP1L Ab enriched 103 centrosomal proteins, while the LUZP1 Ab enriched only 5, suggesting that FILI1L binds DAPK3 at the centrosome (**Fig. 3C** and **Table S2**). To cross-validate that FILIP1L interacts with the centrosome, we performed co-IP/MS using a specific Ab recognizing PCM1, a critical centrosomal scaffold protein (**Table S2**). Comparing proteins that co-enriched with LUZP1 and FILIP1L with the PCM1 co-IP validated that the interactomes of PCM1 and FILIP1L overlapped by 41% (*N* = 42), while LUZP1 overlapped only by 3.8% (*N* = 5); this suggested that primarily FILIP1L localized to the centrosome in mesenchymal-like HCC cells. To confirm that DAPK1, DAPK3, and FILIP1L co-localize at the centrosome, we next performed confocal IF imaging in SNU387 cells. Staining the centrosome with a γ-tubulin Ab showed distinct puncta in the perinuclear region, concordant with typical centrosome localization (**Fig. 3D**). The DAPK3 Ab showed nuclear staining, concordant with its nuclear localization, and stained the same perinuclear puncta as the γ-tubulin Ab; merging both images confirmed co-localization (**Fig 3D**). We also co-stained γ-tubulin and DAPK1, which showed the same pattern of colocalization (**Fig. 3E**), and we co-stained γ-tubulin, DAPK3, and FILIP1L, which likewise showed co-localization at the centrosome (**Fig. 3F**). Collectively, our Co-IP/MS and confocal IF results added evidence that DAPK1, DAPK3, and FILIP1L form a complex at the centrosome. While DAPK3 has been shown to promote cancer cell migration, DAPK1 was shown to mainly suppress migration;^59^ this led us to probe if centrosome-localized DAPK1 was in its active or inactive form. We co-stained γ-tubulin and pDAPK1-S308, a phosphosite that marks inactive DAPK1, in SNU387 cells; this showed, like DAPK1 protein staining, that inactive pDAPK1-S308 localized to the centrosome (**Fig. S3A**). Next, to determine if the components of the DAPK1-DAPK3-FILIP1L complex co-operate to control the same cellular phenotype, we knocked down single complex components and *DAPK1*-*DAPK3* combined in SNU387 cells using transient siRNAi for 3 days, followed by wound scratch and trans-well migration assays (**Fig. 3G** and **S3B**). We validated knockdown by immunoblotting, and an ATP-based viability assay showed that only the *DAPK1*-*DAPK3* double knockdown reduced viability by ∼25% (**Fig. S3C** and **S3D**). Like HS-94 and HS-148, *DAPK3* RNAi strongly inhibited wound closure, as did *DAPK1* and *FILIP1L* knockdown, and *DAPK1*-*DAPK3* double knockdown completely blocked wound closure (**Fig. 3G**). Similarly, knockdown of complex components significantly reduced trans-well invasion (**Fig. S3B**). These results added evidence that the DAPK1-DAPK3-FILIP1L complex forms a functional unit that controls directed HCC cell migration and invasion. Hypothesizing that FILIP1L recruits DAPK3 to the centrosome by means of its leucin zipper domain, and potentially co-recruits DAPK1, we knocked down FILIP1L in SNU387 cells using siRNAi for 3 days and performed confocal IF imaging staining for DAPK1, DAPK3, and γ-tubulin; this showed that FILIP1L knockdown displaced DAPK3 from the centrosome, while DAPK1 was retained at the centrosome (**Fig. 3H**). In contrast, DAPK1 knockdown also abrogated DAPK3 localization to the centrosome (**Fig. S3E**). This suggested that FILIP1L acts as a scaffold protein that recruits DAPK3 to the centrosome, and that inactive pDAPK1-S308 acts as a co-scaffold to FILIP1L.

### 4. The DAPK1-DAPK3-FILIP1L complex controls positioning of the centrosome during directed cell migration through a phosphorylation-dependent mechanism

Positioning of the centrosome towards the leading edge is critical for directed cell migration and cancer cell invasion.^8, 60–63^ Because inhibiting DAPK1-DAPK3-FILIP1L complex components had a strong effect on directed cell migration and invasion, and the complex localizes to the centrosome, we speculated that the complex controls centrosome positioning during the EMPS and directed cell migration.^8^ To quantify if inhibition of the complex alters alignment of the nucleus-centrosome-leading edge axis in migrating cells, we performed a wound scratch assay in SNU387 cells, followed by confocal IF imaging. We then measured the angle α° between the nucleus-scratch axis and the nucleus-centrosome axis (**Fig. 4A**). In cells that can position the centrosome in a way that is conductive of directed migration, α° should be close to zero, and when centrosome repositioning is inhibited, it should be randomly positioned in the peri-nuclear region, like in cells that were not wounded, resulting in a α° anywhere between 0 and 180°. Treating SNU387 cells with the dual DAPK1-DAPK3 inhibitors HS-94 and HS-149 followed by wounding and staining the nucleus and γ-tubulin revealed that cells were less able to re-position the centrosome towards the wound (**Fig. 4B** and **4C**). Similarly, siRNAi-mediated knockdown of *DAPK1*, *DAPK3*, and *FILIP1L*, and double-knockdown of *DAPK1*-*DAPK3*, for 3 days prevented centrosome re-positioning towards the wound (**Fig. 4C**). Concordantly, staining SNU387 cells with an acetylated α-tubulin Ab to visualize the microtubule network revealed that pharmacological DAPK1-DAPK3 inhibition prevented microtubule polarization toward the direction of migration (**Fig. 4D**). This suggested that the DAPK1-DAPK3-FILIP1L complex is critical for centrosome re-positioning and microtubule polarization during directed mesenchymal-like HCC cell migration. Next, to determine if the DAPK1-DAPK3-FILIP1L complex controls centrosome positioning through a phosphorylation dependent mechanism, we treated the mesenchymal-like HCC cell lines SNU387, SNU449, and SNU761 with the DAPK1-DAPK3 inhibitors HS-94 and HS-148 for 3 days, followed by MS-based deep global phosphoproteome profiling; this quantified 16,833 unique phosphosites at a site localization FDR = 0.1 (**Table S3**). Differential expression analysis (DEA) between inhibitor vs. vehicle treated cells showed that 568 sites consistently decreased in abundance, likely representing phosphorylation events downstream of DAPK1 and DAPK3. Filtering for phosphoproteins that have been shown to localize to the centrosome previously (GOCC ‘Centrosome’), revealed that 54 sites on 37 proteins were reduced in phosphorylation. Network analysis using STRING 12.0 revealed that these phosphoproteins belonged to several functional clusters, including a large cluster controlling cell division, a cluster controlling microtubule nucleation, a smaller cluster controlling RhoV signaling, and a cluster controlling centriolar subdistal appendages (**Fig. 4E**).^64^ Predicting the consensus motifs contained in the 54 DAPK1-DAPK3 regulated centrosomal phosphoproteins using The Kinase Library identified eight sites that scored positively for the DAPK1-DAPK3 consensus sequences across different clusters (R-R/K-X-S/T, **Fig. 4E** and **Table S3**).^65^ Two of these sites were located on CEP170 (S1160) and CEP131 (S78), which control microtubule tethering to the centrosome and cell morphology.^66–68^ CEP170-S1160 is an uncharacterized site within the C-terminal domain that mediates centrosome localization,^67^ while CEP131-S78 has been previously characterized as a site that can be phosphorylated by PLK4 and MK2 (MAPKAPK2), and which controls the dynamics of centriolar satellites in response to stress signaling.^69, 70^ Collectively, our phosphoproteomic data suggested that the DAPK1-DAPK3-FILIP1L complex promoted centrosome dynamics during directed mesenchymal cell migration by affecting multiple substrates involved in centrosome dynamics, including CEP131 and CEP170.

**Figure 4.**
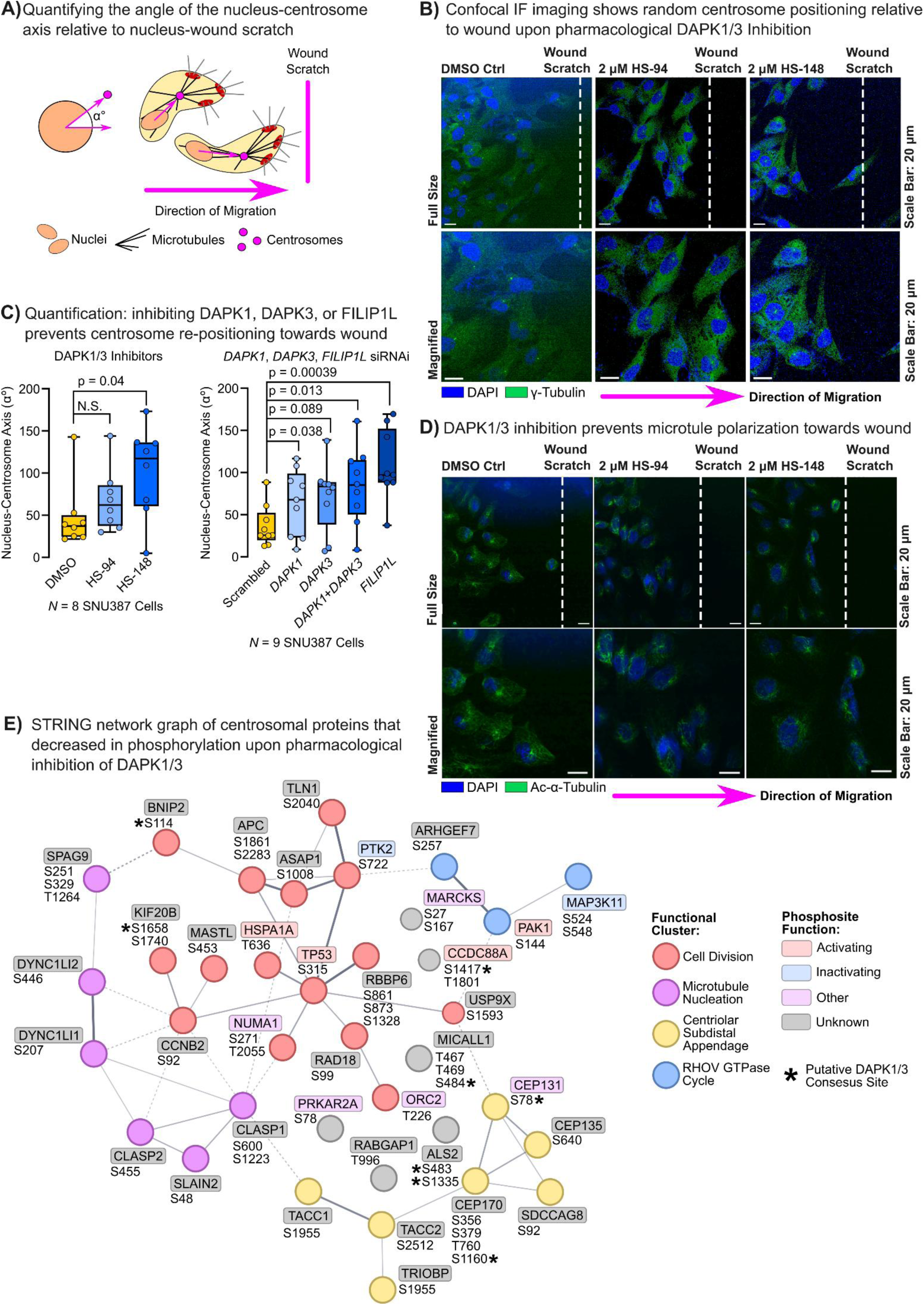
The DAPK1-DAPK3-FILIP1L complex controls centrosome positioning and microtubule polarization in a phosphorylation-dependent manner. **(A)** Schematic showing our strategy to quantify deviation of the nucleus-centrosome-wound scratch axis (angle α°) in mesenchymal-like HCC cells that perform directed cell migration. **(B)** Confocal IF images of SNU387 cells that have been pre-treated with 2 µM of either of the dual DAPK1-DAPK3 inhibitors HS-94 and HS-148 for 3 days followed by scratching of a wound; Images were taken 6 h after scratching. Stained were the nucleus (DAPI) and the centrosome (γ-tubulin). **(C)** Same experimental setup as in (B) but stained were the nucleus (DAPI) and the cellular microtubule network (Ac-α-tubulin). **(D)** Quantification of angles α° as determined by confocal IF imaging in SNU387 cells. Cells were pretreated with DAPK1-DAPK3 inhibitor or siRNAs targeting DAPK1, DAPK3, FILIP1L, or both DAPK1-DAPK3 for 72 h; controls were DMSO (inhibitors) or a scramble siRNA sequence. Statistics: two-sided two-sample Student’s T-test, p < 0.05. **(E)** Results of MS-based global phosphoproteomics analysis of SNU387, SNU449, and SNU761 cells that were treated with the either of the DAPK1-DAPK3 inhibitors for 3 d. Shown is a String 12.0 network of phosphoproteins (GOCC ‘Centrosome’) that significantly diseased in phosphorylation in response to the DAPK1-DAPK3 inhibitors. Edges represent confidence and k-means clustering was performed allowing four clusters. Statistics: two-sided two-sample Student’s T-test, p < 0.05 for both inhibitors in the same cell line.

## DISCUSSION

To migrate and invade tissues, carcinoma cells typically must switch from the apical-basal polarity of nonmotile epithelial-like cells to the rear-front polarity of motile mesenchymal-like cells.^3^ Knowledge of the pathways that control this EMPS is widely dispersed, and systematic studies to determine the kinase signaling networks that underly the EMPS are lacking. We integrated quantitative MS-based kinome profiling with chemical genetic and QPI phenotypic screening and wound healing assays to identify the kinases that promote polarity switching and directed cell migration, an approach that we dubbed morphokin-MS.^37–41^ Treating four distinct epithelial-like HCC cell lines with a morphogenic, EMT-inducing cytokine and Ab cocktail, followed by MS-based kinome profiling showed that kinome reprogramming is heterogeneous among different cell lines, but also that a conserved kinase network emerged that was frequently induced during the EMPS (**Fig. 2A** and **2B**, **Table S1**). Chemical genetic screening with selective KIs, followed by QPI analysis of cell morphology and wound healing assay, determined those kinases that actively drive the HCC cell EMPS and directed mesenchymal-like cell migration. Several of the EMPS-associated kinases that we identified have been linked to cell polarity independently from one another previously, including DDR1 and WNK1, while the function of our other screening hits like EPHB2 and MAP4K4 in the EMPS remains obscure.^71, 72^ Our MS-based kinome profiling of 17 HCC patient tumors with diverse disease etiologies showed that 5/12 EMPS-associated kinases were frequently increased in abundance in tumors (*N* ≥ 8 cases), which correlated with poor patient outcomes. Tumor kinome profiling also revealed dozens of additional kinases that were highly abundant in tumors, and that may promote aggressive tumor traits and present novel HCC drug targets (**Fig. S2A** and **Table S1**). Collectively, our comprehensive kinome profiling data will serve the cancer research community as a valuable resource for mechanistic studies into cancer cell polarity and invasiveness, and HCC drug discovery.

Studying DAPK3’s function in the EMPS in greater detail, we found that its pharmacological inhibition and knockdown reversed the EMPS and blocked directed cell migration. We showed that DAPK3 physically interacted with DAPK1 and FILIP1L at the centrosome to promote its re-positioning towards the leading edge of migrating mesenchyme-like HCC cells; centrosome repositioning is a perquisite for directed mesenchymal-like cell migration.^8^ DAPK3, DAPK1, and FILIP1L have been linked to cancer cell migration and invasion independently from one another before. DAPK3 was initially characterized as an apoptosis-inducing kinase and was later found to control, among others, actin dynamics and focal adhesion reorganization, and cancer cell invasion and migration.^53, 73–75^ Importantly, *DAPK3* knockdown was also shown to disrupt fibroblast morphology and directed cell migration; this is concordant with our data suggesting that DAPK3 partakes in specialized mesenchymal cell polarity programs.^76^ DAPK1 has, with few exceptions, been shown to suppress cancer cell migration and mainly acts as a tumor suppressor,^59^ and FILIP1L’s function in EMT-like transitions and cancer cell invasions is context dependent.^56, 58, 77–79^ Here we present a mechanistic link between DAPK1, DAPK3, and FILIP1L, which physically interact at the centrosome and form a functional complex that promotes front-rear polarity, directed migration, and invasion of mesenchymal-like HCC cells by promoting centrosome re-positioning and microtubule polarization towards the cell’s leading edge. Specifically, our data suggested that FILIP1L and inactive, S308 phosphorylated, DAPK1 act as scaffold proteins that co-recruit DAPK3 to the centrosome. FILIP1L does so likely through heterodimerization of the leucine zipper domains of DAPK3 and FIPLIP1L, and DAPK1 has also been proposed to have scaffold functions previously.^80, 81^ LUZP1, another leucine zipper protein identified in our kinobead AP-MS and co-IP/MS studies has been shown to heterodimerize with DAPK3 previously; it was shown that the leucine zippers were essential to mediate DAPK3 phosphorylation of tight junction proteins and cell migration.^53^ This observation, and our finding that the leucine zipper protein FILIP1L recruited DAPK3 to the centrosome to control its dynamics, led us to speculate that FILIP1L and LUZP1, and possibly other leucine zipper proteins, act as zipper kinase-anchoring proteins (ZipKAPs) that help assemble DAPK3 signaling microdomains throughout the cell, similar to A-kinase anchoring proteins (AKAPs).^82^ Our global phosphoproteomics data from mesenchymal-like HCC cells treated with DAPK1-DAPK3 inhibitors supported that once recruited to the centrosome DAPK3 either directly, or *via* intermediate kinases like PAK1, phosphorylates proteins involved in centrosome and microtubule dynamics, including proteins involved in microtubule anchoring to the centrosome (e.g., CEP135 and CEP170) and nucleus (TACC2; **Fig. 4E**). Collectively, applying our morphokin-MS approach we have discovered a DAPK1-DAPK3-FILIP1L complex that constitutes a centrosomal signaling micro domain, controlling centrosome dynamics and positioning during directed mesenchymal-like carcinoma cell migration.

Precise organelle positioning within cells is fundamental to all aspects of cellular and organismal physiology, and is frequently deregulated in cancers, promoting cancer hallmarks like proliferation and invasion.^83, 84^ How kinase signaling controls organelle positioning during directed cancer cell migration and invasion, however, is incompletely understood. The few known examples include the serine/threonine kinases STK24, STK25, and STK26 that control Golgi positioning during the directed cell migration,^85, 86^ AMPK, which controls mitochondrial positioning to the leading edge in invading ovarian cancer cells,^87^ MRCK (CDC42BPA/B/G), which controls nuclear movement and centrosome polarization during directed cell migration,^88^ and AURKA, which has been shown to also control centrosome positioning and microtubule polarization during cell migration.^89^ Our systematic morphokin-MS study identified multiple additional kinases that are activated during the EMPS and may control additional kinase pathways that control organelle positioning. NUAK1, for instance, was strongly induced during the EMPS in all HCC cell lines; NUAK1 has been shown to control mitochondrion positioning in neurons,^90, 91^ and one could speculate that NUAK1 can also control mitochondrion positions to leading edge of migrating cancer cells to control its biogenetics.

In summary, we presented our morphokin-MS approach that broadly identifies kinase networks controlling different aspects of cell morphology and polarity, and we discovered a signaling microdomain that controls centrosome dynamics during mesenchymal-like cell migration. Our comprehensive kinome profiling data from HCC cell lines and patient tissues will serve the cancer and cell biology research communities as an invaluable resource.

## ACKNOWLEDGMENTS

This work was supported by grants from the National Institutes of Health issued under the award numbers 1R35GM150766 (M.G.) and P30CA042014 (M.G.). The content is solely the responsibility of the authors and does not necessarily represent the official views of the National Institutes of Health. T.A.R was supported by the Susan Cooper Jones Endowed Fellowship in Cancer Research and HCI Gastrointestinal Cancers Center trainee grant. A.M.C was supported by the Utah CTSI STARS T32 Training Program fellowship. We thank Dr. Christopher Reilly, Dr. Cassandra Rice, and Erin Romero (University of Utah, Department of Pharmacology & Toxicology) for providing training and access to the ImageXpress Pico Automated Cell Imaging System. We thank Dr. Timothy Haystead for kindly providing the DAPK inhibitors HS-94 and HS-148. We thank the members of the HSC Cell Imaging Core (University of Utah) for providing training and access to the Nikon Spinning Disk Confocal Microscope.

## AUTHOR CONTRIBUTIONS

Conceptualization, M.G. and T.A.R.; Methodology, M.G. and T.A.R.; Investigation, M.G., T.A.R., K.W., P.J., K.W., A.M.C., R.Z., L.H.L., J.Y., R.G.Z, A.T, R.J.T, and P.S.; Formal Analysis, M.G., T.A.R., K.E., R.J.T.; Writing – Original Draft, M.G. and T.A.R.; Writing – Review and Editing, M.G., T.A.R.; Funding Acquisition, M.G and T.A.R, and A.M.C.

## RESOURCE AVAILABILITY

### Lead Contact

Martin Golkowski, Department of Pharmacology & Toxicology, Huntsman Cancer Institute, University of Utah, Salt Lake City, UT 84112, USA

### Materials Availability

As lead contact, Martin Golkowski is responsible for all reagent and resource requests. Please contact Martin Golkowski at with requests and inquiries.

### Data and Code Availability

Bruker MS output files, a detailed list of instrument settings, and DIA-NN output files generated by this study can be requested directly from the principal investigator. This study did not generate new code.

## MATERIALS AND METHODS

### Cell lines and culture conditions

The HCC cell lines PLC/PRF/5, HepG2, Hep3B2.1-7, SNU387, SNU761, and SNU449 were purchased from the American Type Culture Collection (ATCC). The HCC cell line HuH-7 was a kind gift by Dr. Mei Y. Koh of the University of Utah, Department of Pharmacology and Toxicology. Cell line identity was confirmed by short tandem repeats (STR) marker analysis. All cells were grown at 37°C under 5% CO_2_, 95% ambient atmosphere.

Fifteen cryo-frozen cell stocks were generated from the original vial from the cell bank (passage 3). Experiments were performed with cells at <10 passages from the original vial. All cell media used were those recommended by the ATCC, except HuH-7, which were cultured in DMEM (Gibco). All media were supplemented with 100x penicillin-streptomycin-glutamine (Thermo Fisher Scientific, Gibco), 10% fetal bovine serum (FBS, Corning), and MycoZap mycoplasma elimination reagent (Lonza). Cells were tested once every week for mycoplasma contamination using the MycoStrip 2.0 mycoplasma detection kit (InvivoGen). Cells were harvested or used for other experiments when reaching 90% confluency, unless noted otherwise.

### EMT induction experiments

Epithelial-like HCC cell lines were grown to the given confluency, i.e., HuH-7 cells to 50%, HepG2 to 75%, PLC/PRF/5 to 90%, and Hep3B2.1-7 to 90%; additionally, HepG2 cells were grown on poly-D-lysine-coated tissue culture plates to promote cell adhesion and suppress apoptosis. Depending on the experiment, cells were either kept in complete growth medium, or rinsed twice with sterile PBS, FBS-free growth medium was added, and 1% final of the StemXVivo EMT Inducing Media Supplement (100X, R&D Systems; EMT mix) was added. Different confluences were used before treatment to account for cell line-specific differences in the kinetics by which the EMT mix induced cell cycle arrest. Control cells were grown in FBS-containing or FBS-free medium with water added as the vehicle control. Cells were placed in the tissue culture incubator for 96 h and then processed as described below.^37, 38^

### Transient siRNAi knockdown

siRNA and scrambled control sequences were transfected using the Lipofectamine RNAiMAX Transfection Reagent (Invitrogen) following the manufacturer’s instructions and cells placed in the tissue culture incubator for 72 h, followed by the respective experiments (see below). siRNA sequences were all Life Technology Stealth RNAi sequences, i.e., DAPK3HSS102646 (Alias DAPK3 siRNA#1; Seq. ACGGCGUUCACUACCUGCACU-CUAA and UUAGAGUGCAGGUAGUGAACGCCGU), DAPK3HSS102647 (Alias DAPK3 siRNA#2; Seq.

GGGAACGAGUUCAAGAACAUCUUCG and CGAAGAUGUUCUUGAACUCGUUCCC), FILIP1LHSS117502 (Alias FILIP1L siRNA#1; seq. GCUAAUGAACGAGACAAAGCUCAAU and AUUGAGCUUUGUCUCGUUCAU-UAGC), FILIP1LHSS117503 (Alias FILIP1L siRNA#2; Seq. GAGGAGCAGUGCAGAGAUCUCAAUA and UAUUGAGAUCUCUGCACUGCUCCUC), DAPK1HSS102643 (Alias DAPK1 siRNA#1, Seq.

GCCCUAGCCAAAGAUUUCAUAAGAA and UUCUUAUGAAAUCUUUGGCUAGGGC) and DAPK1HSS102644 (Alias DAPK1 siRNA#2; Seq. CCACGUCAAUCCAUGUUGUUGUCUU and AAGA-CAACAACAUGGAUUGACGUGG).

### Kinase inhibitor treatment experiments

Cells were grown for 72 h in the presence of 2 µM of the KIs in DMSO at a 0.1% (v/v) final concentration of DMSO, which was also the vehicle control. Thereafter, cells were used for experiments as indicated below. KIs and their main targets were HTH-01-015 (NUAK1; MedChemExpress, MCE, Cat# HY-15802) WZ4003 (NUAK1 and NUAK2; MCE, Cat# HY-12334), HS-94 and HS-148 (DAPK1 and DAPK3, kind gifts from Dr. Timothy Haystead of Duke University, Department of Pharmacology & Cancer Biology),^92^ BMPR2-IN-1 (BMPR2, MCE, Cat# HY-154970), DDR1-IN-1 (DDR1, MCE, Cat# HY-13979), MAP4K4-IN-3 (MAP4K4, MCE, Cat# HY-125012), ALW-II-49-7 (EPHB2, Cat# HY-18833), SLK/STK10-IN-1 (SLK and STK10, MCE, HY-132868), saracatinib (SFKs, MCE, HY-10234).

### Trans well migration and invasion assay

Trans well migration and invasion assay were performed as described previously,^37^ with the following minor modifications. For invasion assays, 6-well trans well inserts (Corning, 24-mm inserts, 8 µm pore size) were coated with Corning Matrigel Matrix according to manufacturer instruction. For experiments using pharmacological inhibitors cells were pre-treated as described in ‘Kinase inhibitor treatment experiments‘ above and KIs were replenished after seeding on the well plate inserts. For siRNAi knockdown experiments, cells were pretreated as described under ‘Transient siRNAi knockdown’ above. For EMT induction experiments with HuH-7 cells 1% of the EMT mix or vehicle were applied at the time of seeding, and cells allowed to migrate for 48h. Paclitaxel was positive control for inhibiting migration and invasion and was applied at 100 nM at the time of seeding on the well plate inserts.

### Wound Scratch Assay

20,000 cells in 50 µl growth medium per well (SNU387 and SNU449) were seeded on 96 Well TC-treated microplates (Corning, Cat#3596) and allowed to adhere overnight. Then, 50 µl additional growth medium was added either containing pharmacological inhibitors (0.1% DMSO final) or siRNAs plus Lipofectamine RNAiMAX Transfection Reagent (Invitrogen) and placed in the tissue culture incubator for 72 h. DMSO vehicle, a scramble siRNA sequence, or transfection reagent alone were the controls. Then, the growth medium was carefully aspirated using a micropipette, cells were rinsed with 100 µL sterile PBS, and 15 µM Cell Tracker Green (CTG) in 50 µL of phenol red-free and FBS-free medium was added to each well and cells placed in in the tissue culture incubator for 30 min. Then, the medium was carefully aspirated, cells rinsed twice with 100 µL sterile PBS and wounds were scratched 5-6 times using an Incucyte WoundMaker (Sartorius). Cells were rinsed with 100 µL sterile PBS, replenished with 100 µL phenol red-free growth medium containing FBS, pharmacological inhibitors in DMSO, DMSO vehicle (0.1% DMSO final) were added, and cells placed in the ImageXpress Pico Automated Cell Imaging System (Molecular devices; 5% CO_2_, 15% O_2_ at 37°C). For image acquisition, the following settings were selected: 10 CTG growth program; FITC stain was selected (green); transmitted light (TL), select “capture first”; 4x magnification; focus was adjusted as needed. Images were captured every 2 h for 48 – 96 h depending on the experiment.

### Immunoblotting

Immunoblotting was performed as previously described.^38^ For EMT induction experiments, antibodies used were albumin rabbit polyclonal antibody (ALB, Cell Signaling Technology, CST, Cat# 4929), transferrin (E7F4T) rabbit monoclonal antibody (TF, CST, Cat# 35293), SMAD3 (C67H9) rabbit monoclonal antibody (CST, Cat# 9523), JunB (C37F9) rabbit monoclonal antibody (CST, Cat# 3753), fibronectin/FN1 (E5H6X) rabbit monoclonal antibody (CST, Cat# 26836), vimentin (D21H3) rabbit monoclonal antibody (VIM, CST, Cat# 5741), betaactin (8H10D10) mouse monoclonal antibody (ACTB, Cell Signaling Technology, Cat# 3700), ZIP-kinase (G-1) mouse monoclonal antibody (DAPK3, Santa Cruz Biotechnology, SCBT, Cat# sc-514331), NUAK1/ARK5 (A-9) mouse monoclonal antibody (SCBT, Cat# sc-271827), EphB2 (D2X2I) rabbit monoclonal antibody (CST, Cat# 83029), HGK rabbit polyclonal antibody (MAP4K4, CST, Cat# 3485), DDR1 (D1G6) rabbit monoclonal antibody (CST, Cat# 5583), LOK rabbit polyclonal antibody (STK10, Bethyl, Cat# A300-400A), BMPR2 (E9U5C) rabbit monoclonal antibody (CST, Cat# 24803), histone H3 (D1H2) rabbit monoclonal antibody (Hist. H3, CST, Cat# 4499), profilin-1 rabbit polyclonal antibody (CST, Cat# 3237), Paxillin (GT7612) mouse monoclonal antibody (Novus Biologicals, NB, Cat# NBP3-13563), DOC1 (D-2) mouse monoclonal antibody (FILIP1L, SCBT, Cat# sc-376472), DAPK1 rabbit polyclonal antibody (CST, Cat# 3008).

### Kinase affinity enrichment, KI competition, and on-bead digestion of proteins

Cell lysate preparation, kinase affinity enrichment, KI competition, and on-bead digestion of proteins were performed as previously described.^37, 38^ For kinome abundance profiling in cell lines, epithelial-like HCC cell lines were cultured on 10 cm dishes and treated with EMT mix as described under ‘EMT induction experiments’ above. For kinome abundance profiling in tissue samples, fresh-frozen tissue samples of 40-50 mg wet weight were added to 500 µL of modified RIPA buffer (50 mM Tris-HCl, 150 mM NaCl, 1% NP-40 (v/v), 0.25% Na-deoxycholate (w/v), 1 mM EDTA, 10 mM NaF, 5% glycerol (v/v), MiliQ water, pH 7.8) in a 5 mL polyethylene tubes and kept on wet ice. Tissues were homogenized using a hand-held homogenizer (VWR 200 Homogenizer), lysates incubated on an end-over-end rotator for 30 min at 4°C, and further processed as described.^37, 38^ For KI kinome selectivity profiling, SNU761 cells were grown and lysates at 4 mg/mL protein content generated as described.^37, 38^ The following KIs were used as the competitors for soluble competition experiments at the given final concentrations: HS-94 and HS-148 (both 2 µM and 200 nM). Final DMSO (vehicle) concentration in all pulldowns was 0.1% (v/v).

### Peptide preparation for global proteome and phosphoproteome analysis

Peptide preparation for global proteome and phosphoproteomic analysis was performed as described,^37, 93^ with the following minor modifications. For experiments with pharmacological inhibitors, the HCC cell lines SNU387, SNU449, and SNU761 were grown on 10 cm tissue culture dishes in complete medium in the presence of 2 µM of the dual DAPK1 and DAPK3 inhibitors HS-94 and HS-148 in DMSO at 0.1% (v/v) final concentration.

For EMT induction experiments, epithelial HCC cells were grown on 10 cm dishes and treated with EMT mix as described in ‘EMT induction experiments’ above. Briefly, cells were rinsed twice with ice-cold PBS, lysed in 8 M urea buffer (100 mM Tris, pH 8.5) containing 5 mM tris(2-carboxyethyl) phosphine (TCEP) and 10 mM 2-chloroacetamide (CAM), and 1 mg aliquots of protein were partially purified using acetone/trichloroacetic acid (TCA) precipitation. Protein was digested using endoproteinase LysC and trypsin, and peptides desalted using Oasis HLB extraction cartridges (30 mg extraction material). 5% of the total peptide was desalted using StageTips and used for global proteome analysis.^37, 94^ Peptides on oasis cartridges were eluted and subjected to Villen and Gygi ‘in tube’ protocol as described,^93^ using 20 µL of a 50% slurry of Fe-NTA resin derived from the High Select Phosphopeptide Enrichment Kit (Thermo) as the phosphopeptide capture reagent. Phosphopeptides were desalted using StageTips.^94^

### LC-MS analysis of peptide and phosphopeptide samples

LC-MS analysis of peptide and phosphopeptide samples was performed as previously descibed,^37^ with a few minor modifications. Briefly, phosphopeptide samples were split into two, and one half of the sample analyzed using data-dependent acquisition (DDA) with top15 selection for peptide library generation, and the other half of the sample was analyzed using data-independent acquisition with the diaPASEF method,^37, 95^ on a Bruker nanoElute 2 – timsTOF Pro 2 nLC-MS system. Peptides were separated using 15 cm long, 150 µm inner diameter PepSep columns packed with 1.5 µm diameter C18 beads (Bruker) in 45 min long LC gradients (5-30% B) at flow rates of 0.5 µL/min. LC solvents were A (0.1% formic acid in LC-MS-grade water) and B (0.1% formic acid in LC-MS-grade acetonitrile).

### Co-immunoprecipitation

Co-IP/MS analyses were performed as previously described, with few minor modifications.^37, 38^ Briefly, controls for co-IP/MS were either an isotypic GFP antibody (FILIP1L Co-IP), or isotypic normal IgG (all other co-IP/MS). Pulldowns were performed in 3-4 biological replicates, depending on the experiment. Antibodies used were DAPK3 (mouse monoclonal ZIP-kinase Antibody (G-1), Santa Cruz Biotechnology, Cat# sc-514331, alias DAPK3-Ab#1, and rabbit polyclonal anti-ZIPK Antibody, Bethyl, Cat# A304-222A-T, alias DAPK3-Ab#2), FILIP1L (mouse monoclonal DOC1 Antibody (D-2), Santa Cruz Biotechnology, Cat# sc-376472), LUZP1 (rabbit polyclonal anti-LUZP1 antibody, Bethyl, Cat# A-304-634A-T), and PCM1 (mouse monoclonal PCM1 antibody (G-6), Santa Cruz Biotechnology, Cat# sc-398365).

### Computation of MS raw files and downstream bioinformatics analysis

Bruker MS raw files were computed as described previously,^37^ with few minor modifications. Briefly, phosphopeptide DDA files were computed using Fragpipe v22 for spectral library generation,^96^ and libraries used to compute phosphopeptide DIA files in DIA-NN v.1.8.1.^97^ DIA-NN output files were loaded in Perseus, log2 transformed, median normalized and imputed. To identify differentially expressed proteomic features (DEA) between cell lines and treatment conditions, we applied either a two-sample Student’s t-test or a one sample Student’s t-test, applying Benjamini-Hochberg (BH) correction for multiple hypothesis testing (FDR ≤ 0.05, discovery mode in kinobead AP-MS experiments and to analyze 2nd gen kiCCA data), as previously descibed.^37^ Alternatively, we applied a simple p ≤ 0.05 (validation mode in kinobead AP-MS and Co-IP/MS experiments).

### Plotting STRING interaction networks

PPI network models were plotted using the STRING web application version 12.0 with the following settings: Edges were scaled with confidence, and all interactions were considered.^64^ Clustering was performed using the k-means method, allowing 4 clusters.

### Gene Set Enrichment Analysis (GSEA)

As previously described, for gene set enrichment analysis (GSEA), we used the ssGSEA2.0 script in R together with the Gene Ontology: Biological Process (GOBP) gene sets and the Molecular Signature Database (MSigDB) database according to the published workflow.^98^ Briefly, to rank gene names, we calculated a compound score using the two sample Student’s t-test log2 MS intensity difference multiplied by the -log10 p-value. The parameters used for GSEA were: sample.norm.type = “none”, weight = 1, statistic = “area.under.RES”, output.score.type = “NES”, nperm = 1e3, min.overlap = 10, correl.type = “z.score”, par = T, spare.cores = 1, export.signat.gct = T, extended.output = T.

### KinMap Plotting and DEA Score

For plotting of kinome dendrograms we used the KinMap web application.^99^ Changes in kinase abundance were shown as differential expression score (DEA) score, which was calculated as Score_DEA_ = Average log2 FC * Average -Log10 p-Value * Frequency Altered.

### Confocal immunofluorescence (IF) imaging

2.5×10⁴ SNU387 cells in 150 µL complete growth media were seeded on 8-well chamber slides (Nunc Lab-Tek II) and allowed to adhere overnight. Then, 150 µl additional growth medium was added either containing pharmacological inhibitors (0.1% DMSO final) or siRNAs plus Lipofectamine RNAiMAX Transfection Reagent (Invitrogen) and placed in the tissue culture incubator for 72 h. Then, the medium was removed carefully, cells were rinsed three times with sterile PBS containing MgCl_2_ and CaCl_2_ (PBS+); wound scratching was performed with a 300 µL pipette tip during the second rinse if needed, and cells were incubated for 3 min in PBS+ after scratching. 300 µL of fresh growth media was added and the wound allowed to close for the indicated times.

Then, cells were rinsed three times with PBS+ and fixed for 20 min with 4% formaldehyde in PBS+ at RT. Cells were rinsed three times with MgCl_2_/CaCl_2_-free PBS (PBS-) and incubated with 300 µL blocking solution (PBS-, 1% BSA, 1% NGS or 1% NDS) for 1h at room temperature on a rocker. Then cells were rinsed three times with PBS-(5 min for second rinse), permeabilized with 200 µl PBS-containing 0.5% Triton X-100 for 20 min at RT on a rocker, and again rinsed three times with PBS- (5 min for second rinse). Cells were then incubated with primary antibodies at the manufacturers recommended dilution in PBS-containing 1% BSA, 1% NGS or 1% NDS for 1h at RT on a rocker. Antibodies used were y-tubulin Ab (rabbit - Alexa Fluor 488 (IgG), 1:200), Ac-α-tubulin (mouse - Alexa Fluor 546 (IgG2a), 1:167, N-cadherin (CDH2), DAPK3 (mouse - Alexa Fluor 546 or Alexa Fluor 647 (IgG1)); FILIP1L (rabbit - Alexa Fluor 488 or Alexa Fluor 549 (IgG)) at 5:1000 dilution as per the supplier; rabbit - Alexa Fluor 488 (IgG) at 1:200, and DAPK1. Cells were rinsed three times with PBS- (5 min for second rinse). Then, cells were Incubated with secondary antibodies as required, i.e., goat anti-rabbit IgG Alexa Fluor 488 (4ug/mL) and goat anti-mouse IgG Alexa Fluor 546 (4ug/mL) in PBS containing 1% BSA along with DAPI (1:1000) and phalloidin Alexa Flour 647 (Invitrogen) if required for 30 min at room temperature on a rocker. Cells were rinsed three times with PBS- (5 min for second rinse). Chamber slides were mounted on a Fluoromount G and sealed using clear nail polish. Slides were then imaged on the Nikon Ring TIRF/Spinning Disk Confocal microscope equipped with the NIS Elements AR 6.02.03 software and utilizing the spinning disk mode and the 60X WI objective for the following channels: DAPI (405 nm), green (488 nm), red (561 nm), and far-red (647). Primary antibodies for staining were N-Cadherin (D4R1H) rabbit monoclonal antibody (Cell Signaling Technology, Cat# 13116), gamma tubulin (5R3N3) rabbit monoclonal antibody (Novus Biologicals, Cat# NBP3-16852), gamma tubulin (GT4511) mouse monoclonal antibody (GeneTex, Cat# GTX629704), acetyl-alpha-tubulin (Lys40) (D20G3) rabbit monoclonal antibody (Cell Signaling Technology, Cat# 5335), DAP Kinase 3 rabbit polyclonal antibody (GeneTex, Cat# GTX22057), DAPK1 rabbit polyclonal antibody (MyBioSource, Cat# MBS5400868), and phospho-DAPK1 (Ser308) rabbit polyclonal antibody (Invitrogen, Cat# PA5-64779). Primary antibodies were either visualized with AlexaFluor-labeled secondary antibodies, or by direct chemical labeling with AlexaFluor Antibody Labeling Kits (Invitrogen).

### Quantitative Phase Imaging (QPI)

For KI screening, 10,000 SNU387 cells in complete growth medium per well were seeded on 96 Well TC-treated microplates (Eppendorf Cat# 0030 730.119) and allowed to adhere in a cell culture incubator overnight. Then cells were treated either with KIs in DMSO at 2 µM final concentration or DMSO (control) for 72 h, followed by QPI analysis for 48 h with acquisition every 40 min using 10X Plan N objective and the Phasefocus Acquire V3.10.2 software. The field of view (0.5 mm x 1.5 mm) with optimum single-cell segmentation capacity was manually selected for each well. KIs and their main targets were HTH-01-015 (NUAK1; MedChemExpress, MCE, Cat# HY-15802) WZ4003 (NUAK1 and NUAK2; MCE, Cat# HY-12334), HS-94 and HS-148 (DAPK1 and DAPK3, Haystead Lab, Duke University), BMPR2-IN-1 (BMPR2, MCE, Cat# HY-154970), DDR1-IN-1 (DDR1, MCE, Cat# HY-13979), MAP4K4-IN-3 (MAP4K4, MCE, Cat# HY-125012), ALW-II-49-7 (EPHB2, Cat# HY-18833), SLK/STK10-IN-1 (SLK and STK10, MCE, HY-132868), and saracatinib (SFKs, MCE, HY-10234). For EMT induction, 20,000 HuH-7 cells were seeded as described above and allowed to adhere overnight. Complete growth medium was removed and cells carefully rinsed with sterile PBS, and FBS-free growth medium containing 1% final of the StemXVivo EMT Inducing Media Supplement (100X, R&D Systems; EMT-mix). Cells were incubated for 24 h, followed by QPI analysis. Cells were imaged by quantitative phase imaging (QPI) every hour for 96 h under environmentally controlled conditions (37 °C, 5% CO₂, humidified) on a Livecyte 2 imaging system (Phasefocus, Sheffield, UK). Time series were analyzed using the Analyse cell analysis toolbox (Phasefocus) and in house pipelines.^100, 101^ Briefly, each frame in the time series was segmented to identify the position and morphology of individual cells, and segmented objects were tracked across frames to generate single-cell trajectories. Analyse was then used to calculate cell area (segmented object area), sphericity (a dimensionless index of how closely the cell’s optical thickness profile approximates that of a sphere, where higher values indicate more rounded cells and lower values more spread cells), random cell migration (displacement of the object centroid in x and y over time), and doubling time (rate of change in object number over time). Cells contacting the image border or tracked for fewer than 5 consecutive frames were excluded. Data represents 3 technical replicates per condition with on average 215 cell being imaged.

### Reaction Biology in vitro kinase assay

For compound testing *in vitro* we used the commercial HotSpot^TM^ assay from Reaction Biology (Malvern, PA), which was performed as previously described.^102^ Briefly, the tested kinases were CSNK2A1 (CK2α), DAPK3 (ZIPK), and DAPK1; the compounds were HS-94 and HS-148 (Haystead Lab, Duke University),^92^ which were applied at ten different concentrations starting at 50 μM in a 3-fold dilution series. ATP concentration was 1 mM.

## REFERENCES

1. Weiss F, Lauffenburger D, Friedl P. Towards targeting of shared mechanisms of cancer metastasis and therapy resistance. Nat Rev Cancer. 2022;22(3):157–73. doi: 10.1038/s41568-021-00427-0. PubMed PMID: 35013601; PMCID: PMC10399972.

2. Anderson RL, Balasas T, Callaghan J, Coombes RC, Evans J, Hall JA, Kinrade S, Jones D, Jones PS, Jones R, Marshall JF, Panico MB, Shaw JA, Steeg PS, Sullivan M, Tong W, Westwell AD, Ritchie JWA, Cancer Research UK, Cancer Therapeutics CRCAMWG. A framework for the development of effective anti-metastatic agents. Nat Rev Clin Oncol. 2019;16(3):185–204. doi: 10.1038/s41571-018-0134-8. PubMed PMID: 30514977; PMCID: PMC7136167.

3. Peglion F, Etienne-Manneville S. Cell polarity changes in cancer initiation and progression. J Cell Biol. 2024;223(1). Epub 20231213. doi: 10.1083/jcb.202308069. PubMed PMID: 38091012; PMCID: PMC10720656.

4. Lee M, Vasioukhin V. Cell polarity and cancer--cell and tissue polarity as a non-canonical tumor suppressor. J Cell Sci. 2008;121(Pt 8):1141–50. doi: 10.1242/jcs.016634. PubMed PMID: 18388309.

5. Lamouille S, Xu J, Derynck R. Molecular mechanisms of epithelial-mesenchymal transition. Nat Rev Mol Cell Biol. 2014;15(3):178–96. doi: 10.1038/nrm3758. PubMed PMID: 24556840; PMCID: PMC4240281.

6. Dongre A, Weinberg RA. New insights into the mechanisms of epithelial-mesenchymal transition and implications for cancer. Nat Rev Mol Cell Biol. 2019;20(2):69–84. doi: 10.1038/s41580-018-0080-4. PubMed PMID: 30459476.

7. Gandalovicova A, Vomastek T, Rosel D, Brabek J. Cell polarity signaling in the plasticity of cancer cell invasiveness. Oncotarget. 2016;7(18):25022–49. doi: 10.18632/oncotarget.7214. PubMed PMID: 26872368; PMCID: PMC5041887.

8. Burute M, Prioux M, Blin G, Truchet S, Letort G, Tseng Q, Bessy T, Lowell S, Young J, Filhol O, Thery M. Polarity Reversal by Centrosome Repositioning Primes Cell Scattering during Epithelial-to-Mesenchymal Transition. Dev Cell. 2017;40(2):168–84. doi: 10.1016/j.devcel.2016.12.004. PubMed PMID: 28041907; PMCID: PMC5497078.

9. Kawata M, Kondo J, Onuma K, Ito Y, Yokoi T, Hamanishi J, Mandai M, Kimura T, Inoue M. Polarity switching of ovarian cancer cell clusters via SRC family kinase is involved in the peritoneal dissemination. Cancer Sci. 2022;113(10):3437–48. Epub 20220728. doi: 10.1111/cas.15493. PubMed PMID: 35848881; PMCID: PMC9530866.

10. Liu J, Guo Y, Zhang R, Xu Y, Luo C, Wang R, Xu S, Wei L. Inhibition of TRPV4 remodels single cell polarity and suppresses the metastasis of hepatocellular carcinoma. Cell Death Dis. 2023;14(6):379. Epub 20230628. doi: 10.1038/s41419-023-05903-z. PubMed PMID: 37369706; PMCID: PMC10300155.

11. Adachi Y, Kimura R, Hirade K, Yanase S, Nishioka Y, Kasuga N, Yamaguchi R, Ebi H. Scribble mislocalization induces adaptive resistance to KRAS G12C inhibitors through feedback activation of MAPK signaling mediated by YAP-induced MRAS. Nat Cancer. 2023;4(6):829–43. Epub 20230605. doi: 10.1038/s43018-023-00575-2. PubMed PMID: 37277529.

12. Manning G, Whyte DB, Martinez R, Hunter T, Sudarsanam S. The protein kinase complement of the human genome. Science. 2002;298(5600):1912–34. doi: 10.1126/science.1075762. PubMed PMID: 12471243.

13. Fabbro D, Cowan-Jacob SW, Moebitz H. Ten things you should know about protein kinases: IUPHAR Review 14. Br J Pharmacol. 2015;172(11):2675–700. doi: 10.1111/bph.13096. PubMed PMID: 25630872; PMCID: PMC4439867.

14. Fleuren ED, Zhang L, Wu J, Daly RJ. The kinome ’at large’ in cancer. Nat Rev Cancer. 2016;16(2):83– 98. doi: 10.1038/nrc.2015.18. PubMed PMID: 26822576.

15. Lemmon MA, Schlessinger J. Cell signaling by receptor tyrosine kinases. Cell. 2010;141(7):1117– 34. doi: 10.1016/j.cell.2010.06.011. PubMed PMID: 20602996; PMCID: PMC2914105.

16. Gerritsen JS, White FM. Phosphoproteomics: a valuable tool for uncovering molecular signaling in cancer cells. Expert Rev Proteomics. 2021;18(8):661–74. doi: 10.1080/14789450.2021.1976152. PubMed PMID: 34468274; PMCID: PMC8628306.

17. Hornbeck PV, Kornhauser JM, Latham V, Murray B, Nandhikonda V, Nord A, Skrzypek E, Wheeler T, Zhang B, Gnad F. 15 years of PhosphoSitePlus(R): integrating post-translationally modified sites, disease variants and isoforms. Nucleic Acids Res. 2019;47(D1):D433–D41. doi: 10.1093/nar/gky1159. PubMed PMID: 30445427; PMCID: PMC6324072.

18. Ferguson FM, Gray NS. Kinase inhibitors: the road ahead. Nat Rev Drug Discov. 2018;17(5):353–77. doi: 10.1038/nrd.2018.21. PubMed PMID: 29545548.

19. Cohen P, Cross D, Janne PA. Kinase drug discovery 20 years after imatinib: progress and future directions. Nat Rev Drug Discov. 2021;20(7):551–69. doi: 10.1038/s41573-021-00195-4. PubMed PMID: 34002056; PMCID: PMC8127496 member of the Scientific Advisory Boards of Mission Therapeutics, Ubiquigent and Biocatalyst International. D.C. is an employee and shareholder of AstraZeneca. P.A.J. has received consulting fees from AstraZeneca, Boehringer-Ingelheim, Pfizer, Roche/Genentech, Takeda Oncology, ACEA Biosciences, Eli Lilly and Company, Araxes Pharma, Ignyta, Mirati Therapeutics, Novartis, LOXO Oncology, Daiichi Sankyo, Sanofi Oncology, Voronoi, SFJ Pharmaceuticals, Biocartis, Novartis Oncology, Nuvalent, Esai, Bayer, Transcenta and Silicon Therapeutics; receives post-marketing royalties from DFCI-owned intellectual property on EGFR mutations licensed to Lab Corp; has sponsored research agreements with AstraZeneca, Daichi-Sankyo, PUMA, Boehringer-Ingelheim, Eli Lilly and Company, Revolution Medicines, and Astellas Pharmaceuticals; and has stock ownership in Gatekeeper Pharmaceuticals.

20. Roskoski R, Jr. Properties of FDA-approved small molecule protein kinase inhibitors: A 2026 update. Pharmacol Res. 2026;224:108107. Epub 20260124. doi: 10.1016/j.phrs.2026.108107. PubMed PMID: 41587611.

21. Hong Y. aPKC: the Kinase that Phosphorylates Cell Polarity. F1000Res. 2018;7. Epub 20180625. doi: 10.12688/f1000research.14427.1. PubMed PMID: 29983916; PMCID: PMC6020718.

22. Wu Y, Griffin EE. Regulation of Cell Polarity by PAR-1/MARK Kinase. Curr Top Dev Biol. 2017;123:365–97. Epub 20161205. doi: 10.1016/bs.ctdb.2016.11.001. PubMed PMID: 28236972; PMCID: PMC5943083.

23. Bilder D. Epithelial polarity and proliferation control: links from the Drosophila neoplastic tumor suppressors. Genes Dev. 2004;18(16):1909–25. doi: 10.1101/gad.1211604. PubMed PMID: 15314019.

24. Martin-Belmonte F, Perez-Moreno M. Epithelial cell polarity, stem cells and cancer. Nat Rev Cancer. 2011;12(1):23–38. Epub 20111215. doi: 10.1038/nrc3169. PubMed PMID: 22169974.

25. Lin WH, Asmann YW, Anastasiadis PZ. Expression of polarity genes in human cancer. Cancer Inform. 2015;14(Suppl 3):15–28. Epub 20150330. doi: 10.4137/CIN.S18964. PubMed PMID: 25991909; PMCID: PMC4390136.

26. Jung HY, Fattet L, Tsai JH, Kajimoto T, Chang Q, Newton AC, Yang J. Apical-basal polarity inhibits epithelial-mesenchymal transition and tumour metastasis by PAR-complex-mediated SNAI1 degradation. Nat Cell Biol. 2019;21(3):359–71. Epub 20190225. doi: 10.1038/s41556-019-0291-8. PubMed PMID: 30804505; PMCID: PMC6546105.

27. Vogel A, Meyer T, Sapisochin G, Salem R, Saborowski A. Hepatocellular carcinoma. Lancet. 2022;400(10360):1345–62. doi: 10.1016/S0140-6736(22)01200-4. PubMed PMID: 36084663.

28. Llovet JM, Kelley RK, Villanueva A, Singal AG, Pikarsky E, Roayaie S, Lencioni R, Koike K, Zucman-Rossi J, Finn RS. Hepatocellular carcinoma. Nat Rev Dis Primers. 2021;7(1):6. doi: 10.1038/s41572-020-00240-3. PubMed PMID: 33479224.

29. Rumgay H, Arnold M, Ferlay J, Lesi O, Cabasag CJ, Vignat J, Laversanne M, McGlynn KA, Soerjomataram I. Global burden of primary liver cancer in 2020 and predictions to 2040. J Hepatol. 2022;77(6):1598–606. doi: 10.1016/j.jhep.2022.08.021. PubMed PMID: 36208844; PMCID: PMC9670241.

30. Ladd AD, Duarte S, Sahin I, Zarrinpar A. Mechanisms of drug resistance in HCC. Hepatology. 2024;79(4):926–40. doi: 10.1097/HEP.0000000000000237. PubMed PMID: 36680397.

31. Uka K, Aikata H, Takaki S, Shirakawa H, Jeong SC, Yamashina K, Hiramatsu A, Kodama H, Takahashi S, Chayama K. Clinical features and prognosis of patients with extrahepatic metastases from hepatocellular carcinoma. World J Gastroenterol. 2007;13(3):414–20. doi: 10.3748/wjg.v13.i3.414. PubMed PMID: 17230611; PMCID: PMC4065897.

32. Wang X, Zhang L, Dong B. Molecular mechanisms in MASLD/MASH-related HCC. Hepatology. 2024. Epub 20240213. doi: 10.1097/HEP.0000000000000786. PubMed PMID: 38349726; PMCID: PMC11323288.

33. Giannelli G, Koudelkova P, Dituri F, Mikulits W. Role of epithelial to mesenchymal transition in hepatocellular carcinoma. J Hepatol. 2016;65(4):798–808. doi: 10.1016/j.jhep.2016.05.007. PubMed PMID: 27212245.

34. van Zijl F, Zulehner G, Petz M, Schneller D, Kornauth C, Hau M, Machat G, Grubinger M, Huber H, Mikulits W. Epithelial-mesenchymal transition in hepatocellular carcinoma. Future Oncol. 2009;5(8):1169–79. doi: 10.2217/fon.09.91. PubMed PMID: 19852728; PMCID: PMC2963061.

35. Zhang X, Wang J, Liang X, Jiang D, Sun Y, Hu C, Hu F, He Y, Sun Y, Zhang J, Ding J, Cai S, Wang Y, Yang S, Yang K. BAP31-ELAVL1-SPINK6 axis induces loss of cell polarity and promotes metastasis in hepatocellular carcinoma. Int J Biol Sci. 2025;21(4):1632–48. Epub 20250203. doi: 10.7150/ijbs.102566. PubMed PMID: 39990675; PMCID: PMC11844287.

36. Kapil S, Sharma BK, Patil M, Elattar S, Yuan J, Hou SX, Kolhe R, Satyanarayana A. The cell polarity protein Scrib functions as a tumor suppressor in liver cancer. Oncotarget. 2017;8(16):26515–31. doi: 10.18632/oncotarget.15713. PubMed PMID: 28460446; PMCID: PMC5432276.

37. Woods K, Chan AM, Rants’o TA, Sapre T, Mastin GE, Maguire KM, Ong SE, Golkowski M. diaPASEF-Powered Chemoproteomics Enables Deep Kinome Interaction Profiling. J Proteome Res. 2025;24(9):4463–77. Epub 20250827. doi: 10.1021/acs.jproteome.5c00109. PubMed PMID: 40862632; PMCID: PMC12393673.

38. Golkowski M, Lius A, Sapre T, Lau HT, Moreno T, Maly DJ, Ong SE. Multiplexed kinase interactome profiling quantifies cellular network activity and plasticity. Mol Cell. 2023;83(5):803–18 e8. doi: 10.1016/j.molcel.2023.01.015. PubMed PMID: 36736316.

39. Golkowski M, Lau HT, Chan M, Kenerson H, Vidadala VN, Shoemaker A, Maly DJ, Yeung RS, Gujral TS, Ong SE. Pharmacoproteomics Identifies Kinase Pathways that Drive the Epithelial-Mesenchymal Transition and Drug Resistance in Hepatocellular Carcinoma. Cell Syst. 2020;11(2):196–207 e7. doi: 10.1016/j.cels.2020.07.006. PubMed PMID: 32755597; PMCID: PMC7484106.

40. Golkowski M, Vidadala VN, Lau HT, Shoemaker A, Shimizu-Albergine M, Beavo J, Maly DJ, Ong SE. Kinobead/LC-MS Phosphokinome Profiling Enables Rapid Analyses of Kinase-Dependent Cell Signaling Networks. J Proteome Res. 2020;19(3):1235–47. doi: 10.1021/acs.jproteome.9b00742. PubMed PMID: 32037842; PMCID: PMC7537592.

41. Golkowski M, Vidadala RS, Lombard CK, Suh HW, Maly DJ, Ong SE. Kinobead and Single-Shot LC-MS Profiling Identifies Selective PKD Inhibitors. J Proteome Res. 2017;16(3):1216–27. doi: 10.1021/acs.jproteome.6b00817. PubMed PMID: 28102076; PMCID: PMC5663466.

42. Zhang Y, Judson RL. Evaluation of holographic imaging cytometer holomonitor M4(R) motility applications. Cytometry A. 2018;93(11):1125–31. Epub 20181021. doi: 10.1002/cyto.a.23635. PubMed PMID: 30343513; PMCID: PMC7819361.

43. Kamlund S, Janicke B, Alm K, Judson-Torres RL, Oredsson S. Quantifying the Rate, Degree, and Heterogeneity of Morphological Change during an Epithelial to Mesenchymal Transition Using Digital Holographic Cytometry. Applied Sciences. 2020;10(14):4726. PubMed PMID: doi:10.3390/app10144726.

44. Zhang Y, Urquijo MA, Zitnay RG, Marks K, Belote RL, Hansen MMK, Ferita M, Neuendorf HM, Liu T, Smith EA, Mehrabad EM, Hejna M, Moustafa TE, Lange D, Hu M, Vand-Rajabpour F, Done A, Becker CA, Lieberman M, Chang M, Lohman BK, Stubben CJ, Reeves MQ, Zhang X, Weinberger LS, VanBrocklin MW, Deacon DC, Grossman D, Spike BT, Lex A, Boyle GM, Kulkarni R, Zangle TA, Judson-Torres RL. A BRN2:MYC transcriptional axis regulates interconversion between therapy-resistant and tumorigenic phenotypes in melanoma. Cell Rep. 2025;44(12):116675. Epub 20251215. doi: 10.1016/j.celrep.2025.116675. PubMed PMID: 41405996; PMCID: PMC12834598.

45. Hejna M, Jorapur A, Song JS, Judson RL. High accuracy label-free classification of single-cell kinetic states from holographic cytometry of human melanoma cells. Sci Rep. 2017;7(1):11943. Epub 20170920. doi: 10.1038/s41598-017-12165-1. PubMed PMID: 28931937; PMCID: PMC5607248.

46. Scheel C, Eaton EN, Li SH, Chaffer CL, Reinhardt F, Kah KJ, Bell G, Guo W, Rubin J, Richardson AL, Weinberg RA. Paracrine and autocrine signals induce and maintain mesenchymal and stem cell states in the breast. Cell. 2011;145(6):926–40. doi: 10.1016/j.cell.2011.04.029. PubMed PMID: 21663795; PMCID: PMC3930331.

47. Zeisberg M, Neilson EG. Biomarkers for epithelial-mesenchymal transitions. J Clin Invest. 2009;119(6):1429–37. Epub 20090601. doi: 10.1172/JCI36183. PubMed PMID: 19487819; PMCID: PMC2689132.

48. Kawakami Y, Okada H, Nio K, Hayashi T, Seki A, Nakagawa H, Yamada S, Iida N, Shimakami T, Takatori H, Honda M, Kaneko S, Yamashita T. Transcription factor JUNB is required for transformation of EpCAM-positive hepatocellular carcinoma (HCC) cells into CD90-positive HCC cells in vitro. Cell Death Dis. 2025;16(1):319. Epub 20250419. doi: 10.1038/s41419-025-07602-3. PubMed PMID: 40253402; PMCID: PMC12009367.

49. Wendt MK, Balanis N, Carlin CR, Schiemann WP. STAT3 and epithelial-mesenchymal transitions in carcinomas. JAKSTAT. 2014;3(1):e28975. Epub 20140429. doi: 10.4161/jkst.28975. PubMed PMID: 24843831; PMCID: PMC4024059.

50. Tiwari N, Tiwari VK, Waldmeier L, Balwierz PJ, Arnold P, Pachkov M, Meyer-Schaller N, Schubeler D, van Nimwegen E, Christofori G. Sox4 is a master regulator of epithelial-mesenchymal transition by controlling Ezh2 expression and epigenetic reprogramming. Cancer Cell. 2013;23(6):768–83. doi: 10.1016/j.ccr.2013.04.020. PubMed PMID: 23764001.

51. Ortiz MA, Mikhailova T, Li X, Porter BA, Bah A, Kotula L. Src family kinases, adaptor proteins and the actin cytoskeleton in epithelial-to-mesenchymal transition. Cell Commun Signal. 2021;19(1):67. Epub 20210630. doi: 10.1186/s12964-021-00750-x. PubMed PMID: 34193161; PMCID: PMC8247114.

52. Uhlen M, Fagerberg L, Hallstrom BM, Lindskog C, Oksvold P, Mardinoglu A, Sivertsson A, Kampf C, Sjostedt E, Asplund A, Olsson I, Edlund K, Lundberg E, Navani S, Szigyarto CA, Odeberg J, Djureinovic D, Takanen JO, Hober S, Alm T, Edqvist PH, Berling H, Tegel H, Mulder J, Rockberg J, Nilsson P, Schwenk JM, Hamsten M, von Feilitzen K, Forsberg M, Persson L, Johansson F, Zwahlen M, von Heijne G, Nielsen J, Ponten F. Proteomics. Tissue-based map of the human proteome. Science. 2015;347(6220):1260419. doi: 10.1126/science.1260419. PubMed PMID: 25613900.

53. Wang J, Tran-Huynh AM, Kim B-J, Chan DW, Holt MV, Fandino D, Yu X, Qi X, Wang J, Zhang W, Wu Y-H, Anurag M, Zhang XHF, Zhang B, Cheng C, Foulds CE, Ellis MJ. Death-associated protein kinase 3 modulates migration and invasion of triple-negative breast cancer cells. PNAS Nexus. 2024;3(9):pgae401. doi: 10.1093/pnasnexus/pgae401.

54. Hyodo T, Asano-Inami E, Ito S, Sugiyama M, Nawa A, Rahman ML, Hasan MN, Mihara Y, Lam VQ, Karnan S, Ota A, Tsuzuki S, Hamaguchi M, Hosokawa Y, Konishi H. Leucine zipper protein 1 (LUZP1) regulates the constriction velocity of the contractile ring during cytokinesis. FEBS J. 2024;291(5):927–44. Epub 20231206. doi: 10.1111/febs.17017. PubMed PMID: 38009294.

55. Goncalves J, Sharma A, Coyaud E, Laurent EMN, Raught B, Pelletier L. LUZP1 and the tumor suppressor EPLIN modulate actin stability to restrict primary cilia formation. J Cell Biol. 2020;219(7). doi: 10.1083/jcb.201908132. PubMed PMID: 32496561; PMCID: PMC7337498.

56. Kwon M, Lee SJ, Wang Y, Rybak Y, Luna A, Reddy S, Adem A, Beaty BT, Condeelis JS, Libutti SK. Filamin A interacting protein 1-like inhibits WNT signaling and MMP expression to suppress cancer cell invasion and metastasis. Int J Cancer. 2014;135(1):48–60. Epub 20140220. doi: 10.1002/ijc.28662. PubMed PMID: 24327474; PMCID: PMC3991758.

57. Bozal-Basterra L, Gonzalez-Santamarta M, Muratore V, Bermejo-Arteagabeitia A, Da Fonseca C, Barroso-Gomila O, Azkargorta M, Iloro I, Pampliega O, Andrade R, Martin-Martin N, Branon TC, Ting AY, Rodriguez JA, Carracedo A, Elortza F, Sutherland JD, Barrio R. LUZP1, a novel regulator of primary cilia and the actin cytoskeleton, is a contributing factor in Townes-Brocks Syndrome. Elife. 2020;9. doi: 10.7554/eLife.55957. PubMed PMID: 32553112; PMCID: PMC7363444.

58. Kwon M, Rubio G, Nolan N, Auteri P, Volmar JA, Adem A, Javidian P, Zhou Z, Verzi MP, Pine SR, Libutti SK. FILIP1L Loss Is a Driver of Aggressive Mucinous Colorectal Adenocarcinoma and Mediates Cytokinesis Defects through PFDN1. Cancer Res. 2021;81(21):5523–39. Epub 20210820. doi: 10.1158/0008-5472.CAN-21-0897. PubMed PMID: 34417201; PMCID: PMC8563430.

59. Zhang X, Assaraf YG, Lin Y. Death-associated protein kinase 1: a double-edged sword in health and disease. Front Immunol. 2025;16:1593394. Epub 20250821. doi: 10.3389/fimmu.2025.1593394. PubMed PMID: 40918124; PMCID: PMC12408332.

60. LoMastro GM, Holland AJ. The Emerging Link between Centrosome Aberrations and Metastasis. Dev Cell. 2019;49(3):325–31. doi: 10.1016/j.devcel.2019.04.002. PubMed PMID: 31063752; PMCID: PMC6506172.

61. Luxton GW, Gundersen GG. Orientation and function of the nuclear-centrosomal axis during cell migration. Curr Opin Cell Biol. 2011;23(5):579–88. Epub 20110830. doi: 10.1016/j.ceb.2011.08.001. PubMed PMID: 21885270; PMCID: PMC3215267.

62. Wakida NM, Botvinick EL, Lin J, Berns MW. An intact centrosome is required for the maintenance of polarization during directional cell migration. PLoS One. 2010;5(12):e15462. Epub 20101223. doi: 10.1371/journal.pone.0015462. PubMed PMID: 21203421; PMCID: PMC3009746.

63. Ueda M, Gräf R, MacWilliams HK, Schliwa M, Euteneuer U. Centrosome positioning and directionality of cell movements. Proc Natl Acad Sci U S A. 1997;94(18):9674–8. doi: 10.1073/pnas.94.18.9674. PubMed PMID: 9275182; PMCID: PMC23248.

64. Szklarczyk D, Kirsch R, Koutrouli M, Nastou K, Mehryary F, Hachilif R, Gable AL, Fang T, Doncheva NT, Pyysalo S, Bork P, Jensen LJ, von Mering C. The STRING database in 2023: protein-protein association networks and functional enrichment analyses for any sequenced genome of interest. Nucleic Acids Res. 2023;51(D1):D638–D46. doi: 10.1093/nar/gkac1000. PubMed PMID: 36370105; PMCID: PMC9825434.

65. Johnson JL, Yaron TM, Huntsman EM, Kerelsky A, Song J, Regev A, Lin TY, Liberatore K, Cizin DM, Cohen BM, Vasan N, Ma Y, Krismer K, Robles JT, van de Kooij B, van Vlimmeren AE, Andree-Busch N, Kaufer NF, Dorovkov MV, Ryazanov AG, Takagi Y, Kastenhuber ER, Goncalves MD, Hopkins BD, Elemento O, Taatjes DJ, Maucuer A, Yamashita A, Degterev A, Uduman M, Lu J, Landry SD, Zhang B, Cossentino I, Linding R, Blenis J, Hornbeck PV, Turk BE, Yaffe MB, Cantley LC. An atlas of substrate specificities for the human serine/threonine kinome. Nature. 2023;613(7945):759–66. doi: 10.1038/s41586-022-05575-3. PubMed PMID: 36631611; PMCID: PMC9876800 and is a founder and receives research support from Petra Pharmaceuticals; is listed as an inventor on a patent (WO2019232403A1, Weill Cornell Medicine) for combination therapy for PI3K-associated disease or disorder, and the identification of therapeutic interventions to improve response to PI3K inhibitors for cancer treatment; is a co-founder and shareholder in Faeth Therapeutics; has equity in and consults for Cell Signaling Technologies, Volastra, Larkspur and 1 Base Pharmaceuticals; and consults for Loxo-Lilly. M.B.Y receives research support from Cardiff Oncology. T.M.Y. is a co-founder and stockholder and is on the board of directors of DESTROKE, an early-stage start-up developing mobile technology for automated clinical stroke detection. J.L.J has received consulting fees from Scorpion Therapeutics and Volastra Therapeutics. O.E. is a founder and equity holder of Volastra Therapeutics and OneThree Biotech; is a member of the scientific advisory board of Owkin, Freenome, Genetic Intelligence, Acuamark and Champions Oncology; and receives research support from Eli Lilly, Janssen and Sanofi. D.J.T. is a member of the scientific advisory board at Dewpoint Therapeutics. A.D. is an equity holder of Denali Therapeutics; and receives research support from Interline Therapeutics. N.V. reports consulting activities for Novartis and is on the scientific advisory board of Heligenics. M.D.G. is a co-founder and shareholder of Faeth Therapeutics, which is developing dietary and pharmacological therapies for cancer; and has received speaking and/or consulting fees from Pfizer, Novartis, Scorpion Therapeutics and Faeth Therapeutics.

66. Huang N, Xia Y, Zhang D, Wang S, Bao Y, He R, Teng J, Chen J. Hierarchical assembly of centriole subdistal appendages via centrosome binding proteins CCDC120 and CCDC68. Nat Commun. 2017;8:15057. doi: 10.1038/ncomms15057. PubMed PMID: 28422092; PMCID: PMC5399293.

67. Guarguaglini G, Duncan PI, Stierhof YD, Holmstrom T, Duensing S, Nigg EA. The forkhead-associated domain protein Cep170 interacts with Polo-like kinase 1 and serves as a marker for mature centrioles. Mol Biol Cell. 2005;16(3):1095–107. Epub 20041222. doi: 10.1091/mbc.e04-10-0939. PubMed PMID: 15616186; PMCID: PMC551476.

68. Uetake Y, Terada Y, Matuliene J, Kuriyama R. Interaction of Cep135 with a p50 dynactin subunit in mammalian centrosomes. Cell Motil Cytoskeleton. 2004;58(1):53–66. doi: 10.1002/cm.10175. PubMed PMID: 14983524.

69. Tollenaere MAX, Villumsen BH, Blasius M, Nielsen JC, Wagner SA, Bartek J, Beli P, Mailand N, Bekker-Jensen S. p38-and MK2-dependent signalling promotes stress-induced centriolar satellite remodelling via 14-3-3-dependent sequestration of CEP131/AZI1. Nat Commun. 2015;6:10075. Epub 20151130. doi: 10.1038/ncomms10075. PubMed PMID: 26616734; PMCID: PMC4674683.

70. Denu RA, Sass MM, Johnson JM, Potts GK, Choudhary A, Coon JJ, Burkard ME. Polo-like kinase 4 maintains centriolar satellite integrity by phosphorylation of centrosomal protein 131 (CEP131). J Biol Chem. 2019;294(16):6531–49. Epub 20190225. doi: 10.1074/jbc.RA118.004867. PubMed PMID: 30804208; PMCID: PMC6484138.

71. Hidalgo-Carcedo C, Hooper S, Chaudhry SI, Williamson P, Harrington K, Leitinger B, Sahai E. Collective cell migration requires suppression of actomyosin at cell-cell contacts mediated by DDR1 and the cell polarity regulators Par3 and Par6. Nat Cell Biol. 2011;13(1):49–58. Epub 20101219. doi: 10.1038/ncb2133. PubMed PMID: 21170030; PMCID: PMC3018349.

72. Hwang IY, Kim JS, Harrison KA, Park C, Shi CS, Kehrl JH. Chemokine-mediated F-actin dynamics, polarity, and migration in B lymphocytes depend on WNK1 signaling. Sci Signal. 2024;17(851):eade1119. Epub 20240827. doi: 10.1126/scisignal.ade1119. PubMed PMID: 39190707; PMCID: PMC11542683.

73. She F, Zhang T, Lee TH. Multifaceted role of zipper-interacting protein kinase beyond cell death: Implication of ZIPK dysregulation in neuronal and vascular injuries. Pharmacol Res. 2025;216:107793. Epub 20250521. doi: 10.1016/j.phrs.2025.107793. PubMed PMID: 40409521.

74. Kake S, Usui T, Ohama T, Yamawaki H, Sato K. Death-associated protein kinase 3 controls the tumor progression of A549 cells through ERK MAPK/c-Myc signaling. Oncol Rep. 2017;37(2):1100– 6. doi: 10.3892/or.2017.5359.

75. Wang J, Kim B-J, Anurag M, Yu X, Qi X, Wang J, Zhang B, Cheng C, Ellis M. Abstract PD5-05: PD5-05 DAPK3 modulates migration and invasion of triple negative breast cancers. Cancer Research. 2023;83(5_Supplement):PD5–05–PD5–. doi: 10.1158/1538-7445.SABCS22-PD5-05.

76. Komatsu S, Ikebe M. ZIP kinase is responsible for the phosphorylation of myosin II and necessary for cell motility in mammalian fibroblasts. J Cell Biol. 2004;165(2):243–54. Epub 20040419. doi: 10.1083/jcb.200309056. PubMed PMID: 15096528; PMCID: PMC2172045.

77. Kwon M, Kim JH, Rybak Y, Luna A, Choi CH, Chung JY, Hewitt SM, Adem A, Tubridy E, Lin J, Libutti SK. Reduced expression of FILIP1L, a novel WNT pathway inhibitor, is associated with poor survival, progression and chemoresistance in ovarian cancer. Oncotarget. 2016;7(47):77052–70. doi: 10.18632/oncotarget.12784. PubMed PMID: 27776341; PMCID: PMC5340232.

78. Jing R, Hu C, Qi T, Yue J, Wang G, Zhang M, Wen C, Pei C, Ma B. FILIP1L-mediated cell apoptosis, epithelial-mesenchymal transition and extracellular matrix synthesis aggravate posterior capsular opacification. Life Sci. 2021;286:120061. Epub 20211016. doi: 10.1016/j.lfs.2021.120061. PubMed PMID: 34666037.

79. Gligorijevic B, Belova E, Jarrah A, Abalakov G, Karami A. EMT and cell cycle control invadopodia and metastasis in breast cancer via Filip1L. Res Sq. 2026. Epub 20260419. doi: 10.21203/rs.3.rs-9349300/v1. PubMed PMID: 42040966; PMCID: PMC13105138.

80. Ivanovska J, Tregubova A, Mahadevan V, Chakilam S, Gandesiri M, Benderska N, Ettle B, Hartmann A, Soder S, Ziesche E, Fischer T, Lautscham L, Fabry B, Segerer G, Gohla A, Schneider-Stock R. Identification of DAPK as a scaffold protein for the LIMK/cofilin complex in TNF-induced apoptosis. Int J Biochem Cell Biol. 2013;45(8):1720–9. Epub 20130520. doi: 10.1016/j.biocel.2013.05.013. PubMed PMID: 23702034.

81. Wu YH, Chou TF, Young L, Hsieh FY, Pan HY, Mo ST, Brown SB, Chen RH, Kimchi A, Lai MZ. Tumor suppressor death-associated protein kinase 1 inhibits necroptosis by p38 MAPK activation. Cell Death Dis. 2020;11(5):305. Epub 20200504. doi: 10.1038/s41419-020-2534-9. PubMed PMID: 32366830; PMCID: PMC7198492.

82. Omar MH, Scott JD. AKAP Signaling Islands: Venues for Precision Pharmacology. Trends Pharmacol Sci. 2020;41(12):933–46. Epub 20201017. doi: 10.1016/j.tips.2020.09.007. PubMed PMID: 33082006; PMCID: PMC7890593.

83. Jerabkova-Roda K, Marwaha R, Das T, Goetz JG. Organelle morphology and positioning orchestrate physiological and disease-associated processes. Curr Opin Cell Biol. 2024;86:102293. Epub 20231213. doi: 10.1016/j.ceb.2023.102293. PubMed PMID: 38096602; PMCID: PMC7616369.

84. Kroll J, Renkawitz J. Principles of organelle positioning in motile and non-motile cells. EMBO Rep. 2024;25(5):2172–87. Epub 20240416. doi: 10.1038/s44319-024-00135-4. PubMed PMID: 38627564; PMCID: PMC11094012.

85. Preisinger C, Short B, De Corte V, Bruyneel E, Haas A, Kopajtich R, Gettemans J, Barr FA. YSK1 is activated by the Golgi matrix protein GM130 and plays a role in cell migration through its substrate 14-3-3zeta. J Cell Biol. 2004;164(7):1009–20. doi: 10.1083/jcb.200310061. PubMed PMID: 15037601; PMCID: PMC2172068.

86. Mardakheh FK, Self A, Marshall CJ. RHO binding to FAM65A regulates Golgi reorientation during cell migration. J Cell Sci. 2016;129(24):4466–79. doi: 10.1242/jcs.198614. PubMed PMID: 27807006; PMCID: PMC5201024.

87. Cunniff B, McKenzie AJ, Heintz NH, Howe AK. AMPK activity regulates trafficking of mitochondria to the leading edge during cell migration and matrix invasion. Mol Biol Cell. 2016;27(17):2662–74. Epub 20160706. doi: 10.1091/mbc.E16-05-0286. PubMed PMID: 27385336; PMCID: PMC5007087.

88. Gomes ER, Jani S, Gundersen GG. Nuclear movement regulated by Cdc42, MRCK, myosin, and actin flow establishes MTOC polarization in migrating cells. Cell. 2005;121(3):451–63. doi: 10.1016/j.cell.2005.02.022. PubMed PMID: 15882626.

89. Mori D, Yamada M, Mimori-Kiyosue Y, Shirai Y, Suzuki A, Ohno S, Saya H, Wynshaw-Boris A, Hirotsune S. An essential role of the aPKC-Aurora A-NDEL1 pathway in neurite elongation by modulation of microtubule dynamics. Nat Cell Biol. 2009;11(9):1057–68. Epub 20090809. doi: 10.1038/ncb1919. PubMed PMID: 19668197.

90. Courchet J, Lewis TL, Jr., Lee S, Courchet V, Liou DY, Aizawa S, Polleux F. Terminal axon branching is regulated by the LKB1-NUAK1 kinase pathway via presynaptic mitochondrial capture. Cell. 2013;153(7):1510–25. doi: 10.1016/j.cell.2013.05.021. PubMed PMID: 23791179; PMCID: PMC3729210.

91. Lanfranchi M, Yandiev S, Meyer-Dilhet G, Ellouze S, Kerkhofs M, Dos Reis R, Garcia A, Blondet C, Amar A, Kneppers A, Polveche H, Plassard D, Foretz M, Viollet B, Sakamoto K, Mounier R, Bourgeois CF, Raineteau O, Goillot E, Courchet J. The AMPK-related kinase NUAK1 controls cortical axons branching by locally modulating mitochondrial metabolic functions. Nat Commun. 2024;15(1):2487. Epub 20240321. doi: 10.1038/s41467-024-46146-6. PubMed PMID: 38514619; PMCID: PMC10958033.

92. Carlson DA, Singer MR, Sutherland C, Redondo C, Alexander LT, Hughes PF, Knapp S, Gurley SB, Sparks MA, MacDonald JA, Haystead TAJ. Targeting Pim Kinases and DAPK3 to Control Hypertension. Cell Chem Biol. 2018;25(10):1195–207 e32. doi: 10.1016/j.chembiol.2018.06.006. PubMed PMID: 30033129; PMCID: PMC6863095.

93. Villen J, Gygi SP. The SCX/IMAC enrichment approach for global phosphorylation analysis by mass spectrometry. Nat Protoc. 2008;3(10):1630–8. doi: 10.1038/nprot.2008.150. PubMed PMID: 18833199; PMCID: PMC2728452.

94. Rappsilber J, Mann M, Ishihama Y. Protocol for micro-purification, enrichment, pre-fractionation and storage of peptides for proteomics using StageTips. Nat Protoc. 2007;2(8):1896–906. doi: 10.1038/nprot.2007.261. PubMed PMID: 17703201.

95. Meier F, Brunner AD, Frank M, Ha A, Bludau I, Voytik E, Kaspar-Schoenefeld S, Lubeck M, Raether O, Bache N, Aebersold R, Collins BC, Rost HL, Mann M. diaPASEF: parallel accumulation-serial fragmentation combined with data-independent acquisition. Nat Methods. 2020;17(12):1229–36. doi: 10.1038/s41592-020-00998-0. PubMed PMID: 33257825.

96. Yu F, Teo GC, Kong AT, Frohlich K, Li GX, Demichev V, Nesvizhskii AI. Analysis of DIA proteomics data using MSFragger-DIA and FragPipe computational platform. Nat Commun. 2023;14(1):4154. doi: 10.1038/s41467-023-39869-5. PubMed PMID: 37438352; PMCID: PMC10338508.

97. Demichev V, Messner CB, Vernardis SI, Lilley KS, Ralser M. DIA-NN: neural networks and interference correction enable deep proteome coverage in high throughput. Nat Methods. 2020;17(1):41–4. doi: 10.1038/s41592-019-0638-x. PubMed PMID: 31768060; PMCID: PMC6949130.

98. Krug K, Mertins P, Zhang B, Hornbeck P, Raju R, Ahmad R, Szucs M, Mundt F, Forestier D, Jane-Valbuena J, Keshishian H, Gillette MA, Tamayo P, Mesirov JP, Jaffe JD, Carr SA, Mani DR. A Curated Resource for Phosphosite-specific Signature Analysis. Mol Cell Proteomics. 2019;18(3):576–93. doi: 10.1074/mcp.TIR118.000943. PubMed PMID: 30563849; PMCID: PMC6398202.

99. Eid S, Turk S, Volkamer A, Rippmann F, Fulle S. KinMap: a web-based tool for interactive navigation through human kinome data. BMC Bioinformatics. 2017;18(1):16. doi: 10.1186/s12859-016-1433-7. PubMed PMID: 28056780; PMCID: PMC5217312.

100. Lange D, Polanco E, Judson-Torres R, Zangle T, Lex A. Loon: Using Exemplars to Visualize Large-Scale Microscopy Data. IEEE Trans Vis Comput Graph. 2022;28(1):248–58. Epub 20211224. doi: 10.1109/TVCG.2021.3114766. PubMed PMID: 34587022.

101. Lange D, Judson-Torres R, Zangle TA, Lex A. Aardvark: Composite Visualizations of Trees, Time-Series, and Images. IEEE Trans Vis Comput Graph. 2025;31(1):1290–300. Epub 20241125. doi: 10.1109/TVCG.2024.3456193. PubMed PMID: 39255114; PMCID: PMC12143745.

102. Anastassiadis T, Deacon SW, Devarajan K, Ma H, Peterson JR. Comprehensive assay of kinase catalytic activity reveals features of kinase inhibitor selectivity. Nat Biotechnol. 2011;29(11):1039– 45. doi: 10.1038/nbt.2017. PubMed PMID: 22037377; PMCID: PMC3230241.

