## Supplemental Figures S1-S3 for "Multi-Modal Kinome Profiling Discovers Mesenchymal-Like Polarity Networks that Underly Directed Hepatocellular Carcinoma Cell Migration"

*Supplementary Figures S1-S3*

**Multi-Modal Kinome Profiling Discovers Mesenchymal-Like Polarity Networks that Underly Directed Hepatocellular Carcinoma Cell Migration**

Thankhoe A. Rants'o,[1,2] Kathryn Woods,[1] Paige Jensen,[1] Katie A. Walker,[1,2] Alexandria M. Chan,[1,2] Kathleen M. Maguire,[1] Rebecca G. Zitnay,[2] Lotfa H. Lovely,[1] Jingshu Yang,[3] Augustine Takyi,[2,4] Paul Stewart,[2,4] Kimberley Evason,[2,5] Robert L. Judson-Torres,[2,6] and Martin Golkowski[1,2,*]

[1] Department of Pharmacology and Toxicology, University of Utah, Salt Lake City, UT 84112, USA

[2] Huntsman Cancer Institute, University of Utah, Salt Lake City, UT 84112, USA

[3] Department of Oncological Sciences, University of Utah, Salt Lake City, UT 84112, USA

[4] Department of Nutrition and Integrative Physiology, University of Utah, Salt Lake City, UT 84112, USA

[5] Department of Pathology, University of Utah, Salt Lake City, UT 84112, USA

[6] Department of Dermatology, University of Utah, Salt Lake City, UT 84112, USA

### Figure S1

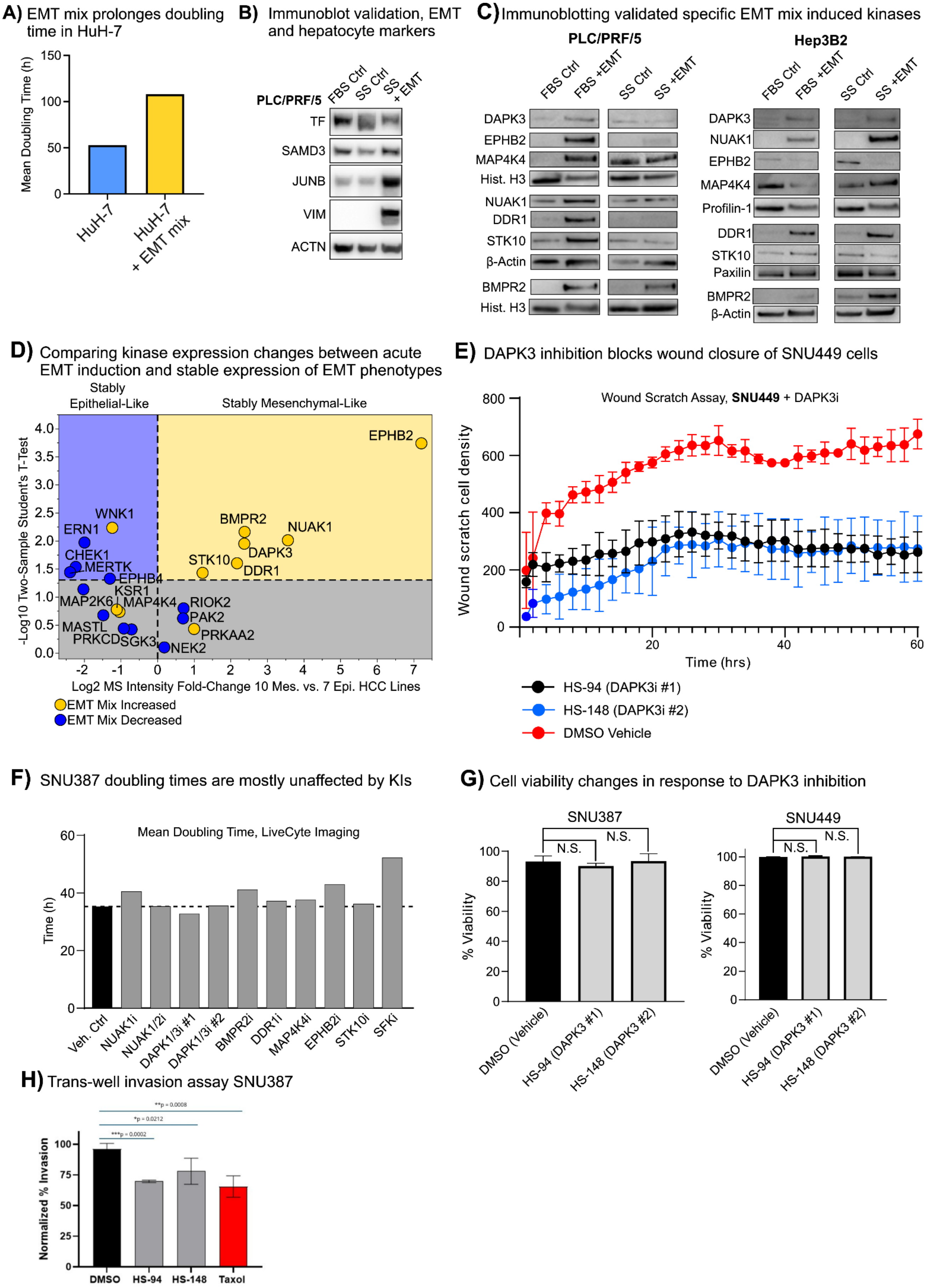
A) EMT mix prolonges doubling time in HuH-7
Mean Doubling Time (h)
150
100
50
0
HuH-7
HuH-7 + EMT mix
B) Immunoblot validation, EMT and hepatocyte markers
PLC/PRF/5
FBS Ctrl
SS Ctrl
SS + EMT
TF
SAMD3
JUNB
VIM
ACTN
C) Immunoblotting validated specific EMT mix induced kinases
PLC/PRF/5
Hep3B2
FBS Ctrl
FBS +EMT
SS Ctrl
SS +EMT
DAPK3
EPHB2
MAP4K4
Hist. H3
NUAK1
DDR1
STK10
β-Actin
BMPR2
Hist. H3
DAPK3
NUAK1
EPHB2
MAP4K4
Profilin-1
DDR1
STK10
Paxilin
BMPR2
β-Actin
D) Comparing kinase expression changes between acute EMT induction and stable expression of EMT phenotypes
Stably Epithelial-Like
Stably Mesenchymal-Like
-Log10 Two-Sample Student's T-Test
Log2 MS Intensity Fold-Change 10 Mes. vs. 7 Epi. HCC Lines
EPHB2
BMPR2
NUAK1
DAPK3
STK10
DDR1
WNK1
ERN1
CHEK1
MERTK
EPHB4
KSR1
MAP2K6
MAP4K4
RIOK2
PAK2
PRKAA2
MASTL
PRKCD
SGK3
NEK2
EMT Mix Increased
EMT Mix Decreased
E) DAPK3 inhibition blocks wound closure of SNU449 cells
Wound Scratch Assay, SNU449 + DAPK3i
Wound scratch cell density
Time (hrs)
HS-94 (DAPK3i #1)
HS-148 (DAPK3i #2)
DMSO Vehicle
F) SNU387 doubling times are mostly unaffected by KIs
Mean Doubling Time, LiveCyte Imaging
Time (h)
Veh. Ctrl
NUAK1i
NUAK1/2i
DAPK1/3i #1
DAPK1/3i #2
BMPR2i
DDR1i
MAP4K4i
EPHB2i
STK10i
SFKi
G) Cell viability changes in response to DAPK3 inhibition
SNU387
SNU449
N.S.
% Viability
DMSO (Vehicle)
HS-94 (DAPK3 #1)
HS-148 (DAPK3 #2)
H) Trans-well invasion assay SNU387
**p = 0.0008
*p = 0.0212
***p = 0.0002
Normalized % Invasion
DMSO
HS-94
HS-148
Taxol

**Figure S1.** Validation of model cell lines treated with EMT, and validation of EMPS-associated kinases. **(A)** Differences in mean doubling time of HuH-7 cells that were treated with EMT mix in serum-free medium compared to serum-free medium alone for 4 days, quantified using the Phasefocus Livecyte QPI system. **(B)** Immunoblot validation of specific EMT markers in HuH-7 and HepG2 cells treated with vehicle in serum-containing medium (FBS ctrl), with vehicle in serum-free medium (serum starvation, SS), and EMT-mix in serum-free medium (SS). Relates to Fig. 1H. **(C)** Immunoblot validation of 7 of the 12 kinases that increased in abundance in response to the EMT mix. Refers to Fig. 2C. **(D)** Differential expression of the 12 EMT mix induced kinases between 7 stably mesenchymal-like and 10 stably epithelial-like HCC cells, as determined by kinobead AP-MS. Statistics: two-tailed two-sample Student's T-test, $p < 0.05$. **(E)** 96-well wound scratch assay of SNU449 cells pre-treated with KIs targeting DAPK3 (HS-94 and HS-148) for 72 h; all KI concentrations were 2 μM. Saracatinib (SFK inhibitor) was positive control. Refers to **Fig. S1E**. Error bars are the S.D. **(F)** Differences in mean doubling time of SNU387 cells that were treated with selective KIs of the 7 validated EMT mix induced kinases for 72 h, quantified using the Phasefocus Livecyte QPI system. All inhibitors were applied at 2 μM concentration. **(G)** Results from a Cell Titer Glo 2.0 assay, quantifying ATP content in SNU387 and SNU449 cells with vehicle (DMSO) or the KIs HS-94 and HS-148 for 72 h; all KI concentrations were 2 μM. *Statistics*: two-tailed two-sample Student's T-test, $p < 0.05$. **(H)** Trans-well invasion assay of HuH-7 cells treated like with DAPK1-DAPK3 inhibitors HS-94 and HS-148 (2 μM), paclitaxel (100 nM), or DMSO vehicle, using FBS as the attractant. Error bars are the standard deviation (S.D.). *Statistics*: two-sided two-sample Student's T-test, $p < 0.05$.

**Figure S2.**

**A)** Kinases that are frequently increased (left) and decreased (right) in abundance in HCC tumors vs. NTL tissues

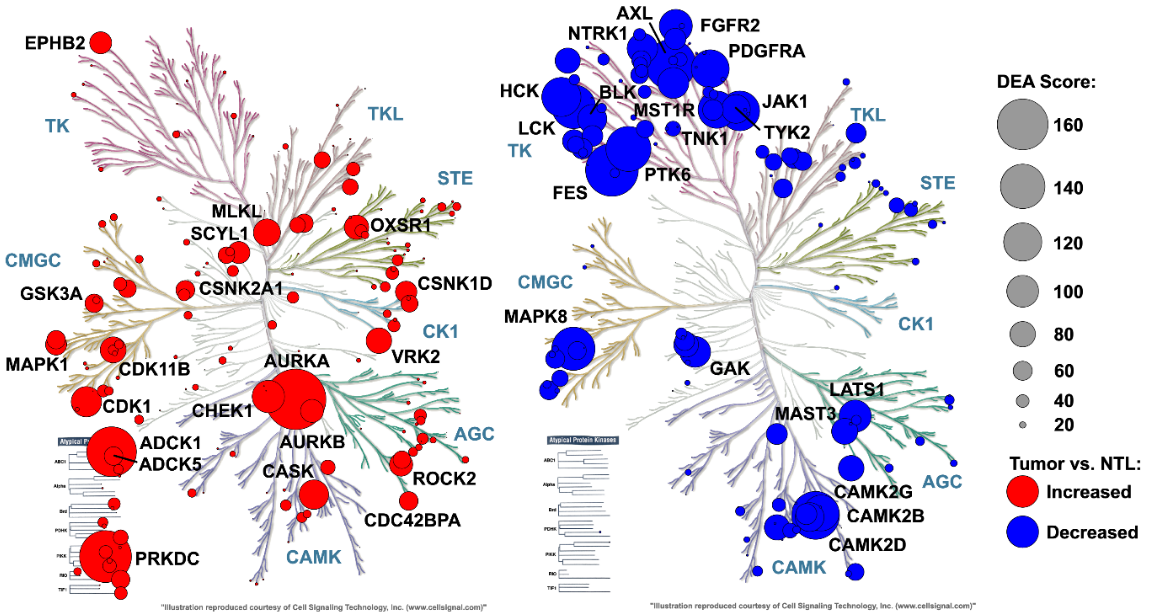


**B)** Kinome profiling of human 17 HCCs and paired NTL tissues reveals frequent activation of kinases linked to EMT-like transitions

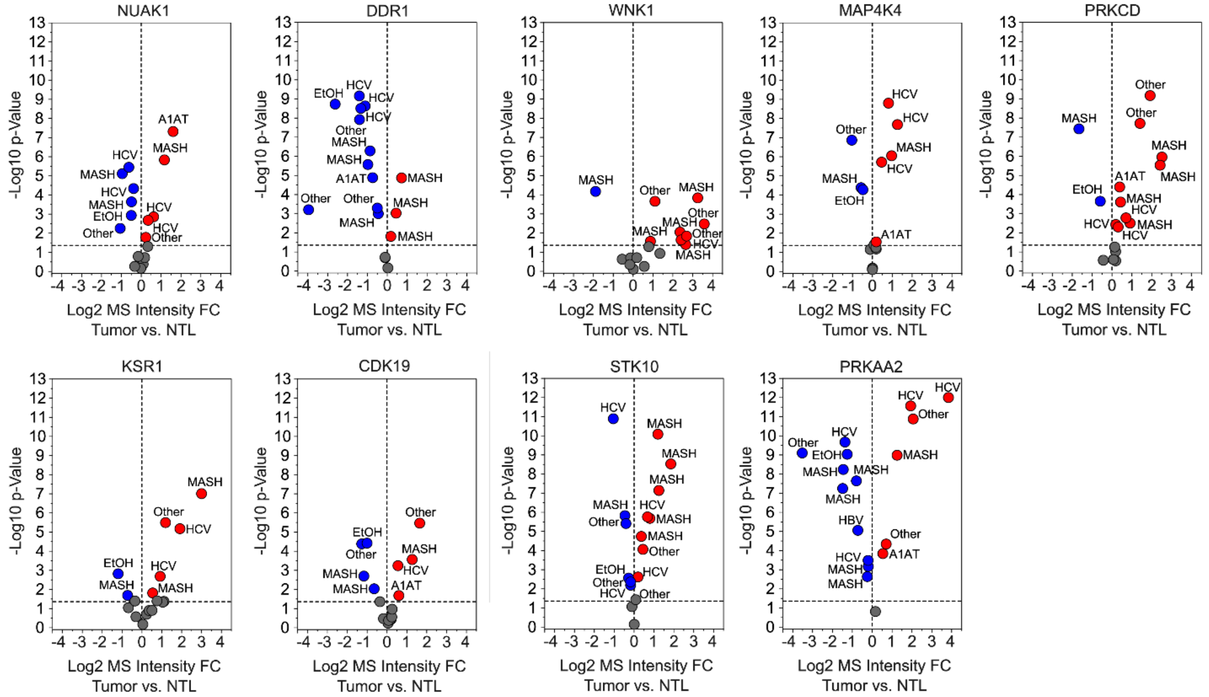


**C)** Epression of kinases linked to EMT-like transitions frequently correlated with poor HCC patient survival

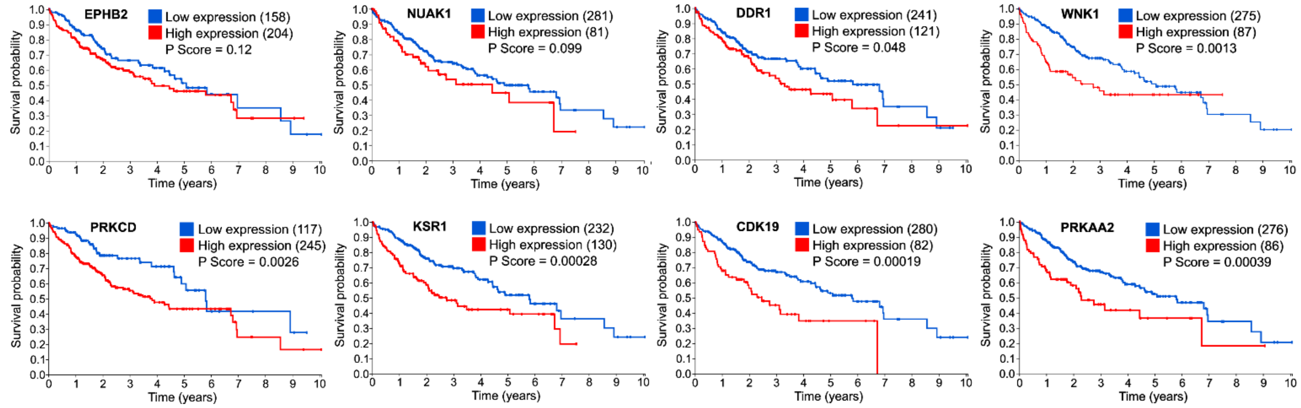

**Figure S2. Kinobead AP-MS profiling of 17 HCC patient tumors and survival analysis for EMPS-associated kinases. (A)** Overlay of the human kinome dendrogram with results from kinobead AP-MS profiling of 17 paired human HCCs and non-tumor liver (NTL) tissue. The DEA score considers the frequency with which specific kinases were significantly increased or decreased between HCC vs. NTL tissues, the log2 fold-change of MS Intensity, and the -log10 p-value from DEA T-test (significance level). *Statistics*: two-sided two-sample Student's T-Test with Benjamini-Hochberg Correction, FDR = 0.05. **(B)** Kinobead AP-MS profiling of 17 HCC patient tumors and paired non-tumor liver (NTL) tissues shows frequently increased abundance of EMP-associated kinases in tumors compared to NTL tissues. Refers to **Fig. 2F**, and **Table S2**. Statistics: see (A). **(C)** Expression of EMPS-associated kinases broadly correlated with shorter patient survival; source was The Protein Atlas. Refers to **Fig. 2G**.

**Figure S3**

**A)** Co-localization of inactive, S-308 phosphorylated DAPK1 with the centrosome

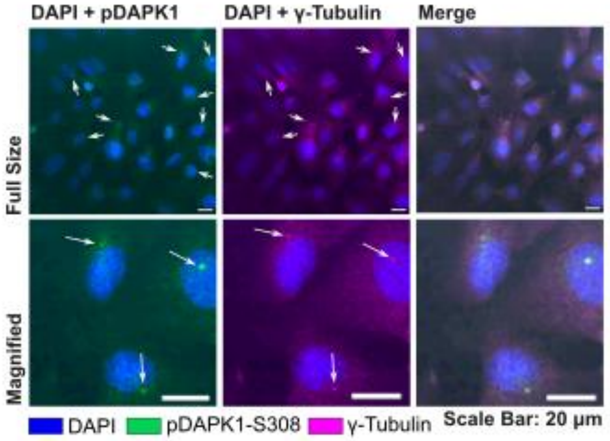


**B)** siRNAi mediated knockdown of *DAPK1-DAPK3-FILIP1L* complex components reduces trans-well invasion

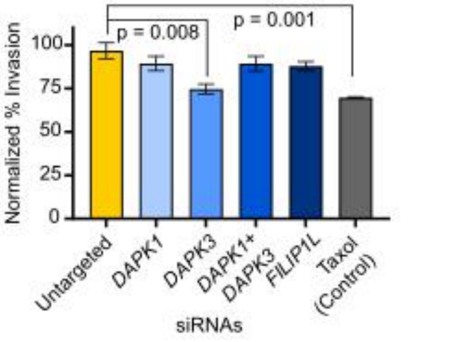


**C)** Immunoblot validation of siRNAi knockdown of *DAPK1*, *DAPK3*, and *FILIP1L* in SNU387

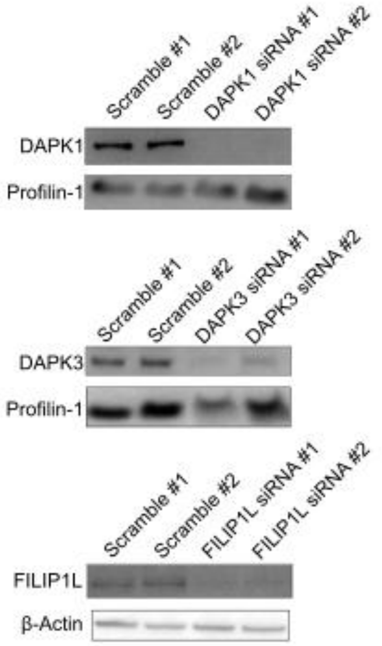


**D)** CellTiter Glo Assay in SNU387 siRNAi lines

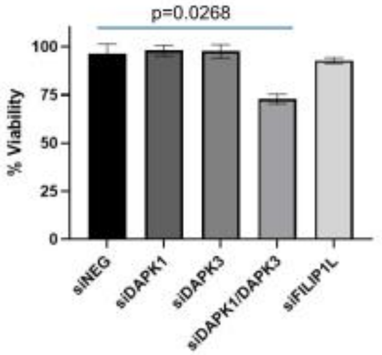


**E)** *DAPK1* knockdown abrogates DAPK3 localization to the centrosome in SNU387 cells

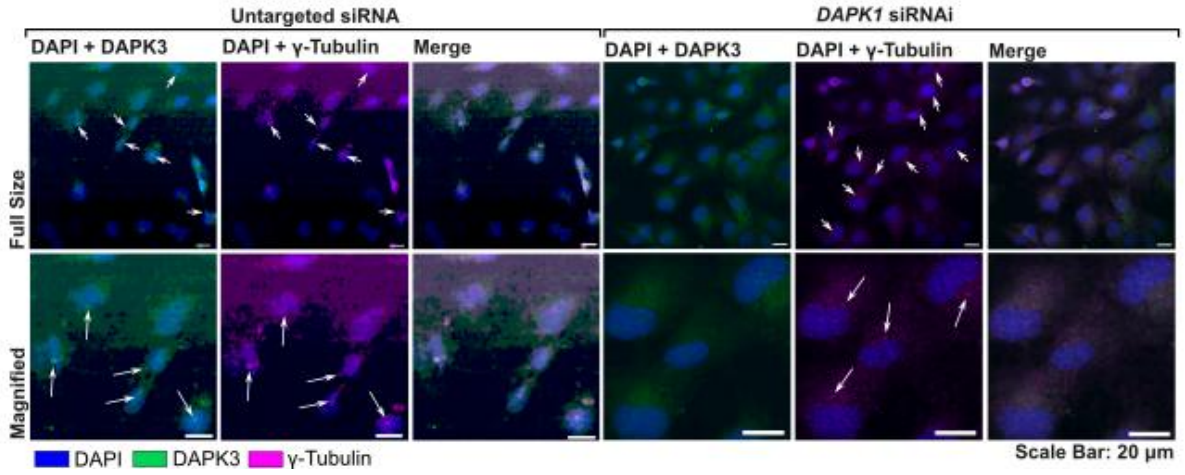

**Figure S3. Validation of DAPK1-DAPK3-FILIP1L colocalization and functional connection.**

**(A)** Co-localization study using confocal IF, staining for the nucleus (DAPI), the centrosome (γ-tubulin), and pDAPK1-S308 in SNU387. Refers to Fig. 3D, 3E, and 3F. **(B)** Trans-well invasion assay of SNU387 cells that were pre-treated with siRNA sequences targeting DAPK1, DAPK3, FILIP1L and DAPK1 + DAPK3, or a scrambled sequence (control) for 3 days. **(C)** Immunoblot validation of *DAPK1*, *DAPK3*, and *FILIP1L* in SNU387 cells that were pre-treated with the respective siRNA sequences or scrambled sequences (control) for 3 days. **(D)** Cell titer Glo assay quantifying ATP content in SNU387 cells that were pretreated with siRNA sequences as described in (B). **(E)** Co-localization study using confocal IF, staining for the nucleus (DAPI), the centrosome (γ-tubulin), and DAPK3 in SNU387 cells that were pretreated with a *DAPK1* siRNA sequence for 3 days. A scrambled, untargeted siRNA sequence was the negative control. Refers to Fig. 3H.
